# Complex interactions between DksA and stress-responsive alternative sigma factors control inorganic polyphosphate accumulation in *Escherichia coli*

**DOI:** 10.1101/2020.03.11.988303

**Authors:** Michael J. Gray

## Abstract

Bacteria synthesize inorganic polyphosphate (polyP) in response to a variety of different stress conditions. PolyP protects bacteria by acting as a protein-stabilizing chaperone, metal chelator, or regulator of protein function, among other mechanisms. However, little is known about how stress signals are transmitted in the cell to lead to increased polyP accumulation. Previous work in the model enterobacterium *Escherichia coli* has indicated that the RNA polymerase-binding regulatory protein DksA is required for polyP synthesis in response to nutrient limitation stress. In this work, I set out to characterize the role of DksA in polyP regulation in more detail. I found that overexpression of DksA increases cellular polyP content (explaining the long-mysterious phenotype of *dksA* overexpression rescuing growth of a *dnaK* mutant at high temperature) and characterized the roles of known functional residues of DksA in this process, finding that binding to RNA polymerase is required, but none of the other functions of DksA appear to be necessary. Transcriptomics revealed genome-wide transcriptional changes upon nutrient limitation, many of which were affected by DksA, and follow-up experiments identified complex interactions between DksA and the stress-sensing alternative sigma factors FliA, RpoN, and RpoE that impact polyP production, indicating that regulation of polyP synthesis is deeply entwined in the multifactorial stress response network of *E. coli*.

**IMPORTANCE:** Inorganic polyphosphate (polyP) is an evolutionarily ancient, widely conserved biopolymer required for stress resistance and pathogenesis in diverse bacteria, but we do not understand how its synthesis is regulated. In this work, I gained new insights into this process by characterizing the role of the transcriptional regulator DksA in polyP regulation in *Escherichia coli* and identifying previously unknown links between polyP synthesis and the stress-responsive alternative sigma factors FliA, RpoN, and RpoE.

## INTRODUCTION

Bacteria, including the model enterobacterium *Escherichia coli*, have sophisticated regulatory systems that allow them to respond to changes in their environments. These systems are essential for survival of stressful or growth-inhibiting conditions, persistence in the environment, and pathogenesis (1–3). In *E. coli*, these include such well-studied systems as the general stress response driven by the alternative sigma factor RpoS (also called *σ*^S^ or *σ*^38^), which upregulates expression of genes involved in resisting starvation, reactive oxygen species, salt, and acid, among others (3, 4) and the stringent response, in which the small molecule alarmones guanosine-5’,3’-tetraphosphate and guanosine-5’,3’-pentaphosphate (collectively referred to as ppGpp) and the RNA polymerase-binding protein DksA coordinately act to slow growth and promote expression of genes involved in adaptation to various kinds of starvation stresses (5–7). Regulation of stress response systems is complex and interconnected, giving bacteria the capacity to survive in rapidly changing environments.

One long-known but poorly-understood element of bacterial stress response is the production of inorganic polyphosphate (polyP), a linear biopolymer of phosphate up to 1000 units in length synthesized by the widely conserved bacterial enzyme polyP kinase (PPK)(8, 9). In response to a variety of stressful changes in conditions, including nutrient limitation, heat, salt, and hypochlorous acid treatment, *E. coli* transiently accumulates large amounts of polyP (10–13). The physiological functions of polyP are not fully understood, but it is known to promote survival under different stress conditions by stabilizing unfolded proteins, chelating toxic metals, acting as a phosphate store, increasing translation fidelity, and regulating the activity of a variety of proteins (12, 14–20). Importantly, in many bacterial pathogens, deleting the gene encoding PPK (*ppk*) results in the loss of the ability to cause disease (21–27), indicating that polyP metabolism may be a potential therapeutic target for use against bacterial infections (28–30).

We know surprisingly little about the mechanism by which polyP synthesis is regulated. Several different regulators have been implicated in modulating polyP production in *E. coli* under different stress conditions, including RpoS (10), the PhoB and PhoU regulators of phosphate transport (10, 13, 31, 32), and the ppGpp synthases RelA and SpoT (10, 13, 33), but no convincing mechanistic model has been developed that explains how any of these systems controls polyP accumulation or why different regulators appear to be required under different conditions. Transcription of the operon containing *ppk* and *ppx*, which encodes the exopolyphosphatase PPX, does not increase upon stress treatment in *E. coli* (11, 34), and while ppGpp potently inhibits degradation of polyP by PPX (33), neither PPX nor ppGpp are necessary for the induction of polyP by nutrient limitation stress (11, 12, 34). However, I have recently reported that the stringent response regulator DksA, which normally works in concert with ppGpp (5), is required for polyP synthesis under those conditions, and this phenotype can be reverted by deletion of the RNA polymerase-binding elongation factor GreA (11), indicating that there is an important role for transcription in polyP control.

In this work, I set out to understand the role of DksA in regulating polyP synthesis in more depth. I found that cellular polyP content can be modulated by changing the amount of DksA expression, and characterized the roles of known functional residues of DksA in this process. I also identified complex interactions between DksA and the alternative sigma factors FliA, RpoN, and RpoE that impact polyP synthesis, gaining new insights into how polyP regulation is linked to the broader stress response network of *E. coli*.

## RESULTS

### Overexpressing DksA enhances stress-induced polyP synthesis

As I have previously reported (11), deletion of *dksA* prevents *E. coli* from accumulating polyP upon nutrient starvation stress (Fig. 1A). In that work, I found that deletion of *greA* in a *dksA* mutant reverts this phenotype and allows polyP to accumulate. However, the *greA*::*cat*^+^ allele used in that and many other studies (35–41) also deletes 109 bp upstream and 177 bp downstream of the *greA* coding sequence (35), potentially disrupting the neighboring *yhbY* gene, which encodes a ribosome assembly factor (42), and the small RNA GraL (43, 44). To confirm that deletion of *greA* is the mutation responsible for allowing a *dksA* mutant to synthesize polyP, I therefore constructed a double mutant strain combining a Δ*dksA1000*::*cat*^+^ mutation (11) with the Δ*greA788*::*kan*^+^ allele from the Keio collection (45), which does not interfere with neighboring gene sequences, and observed that this strain synthesized the same amount of polyP as the wild-type after starvation stress (Fig. 1A). This confirmed that neither *yhbY* nor GraL is involved in regulation of polyP synthesis under these conditions. Interestingly, I also found that overexpressing *dksA* from a strong arabinose-inducible promoter (46) led to accumulation of twice as much polyP as was found in the wild-type strain (Fig. 1B), showing that cellular polyP levels can be tuned by modulating *dksA* expression, although under normal growth conditions the concentration of DksA in the cell is thought to be relatively stable (35). Overexpression of TraR, a protein from the F plasmid that mimics the transcriptional effects of DksA and ppGpp (47, 48), had a similar effect (Fig. 1C), complementing polyP synthesis in a *dksA* mutant and increasing polyP synthesis above wild-type levels in both wild-type and *dksA greA* mutant strains.

**FIG 1.**
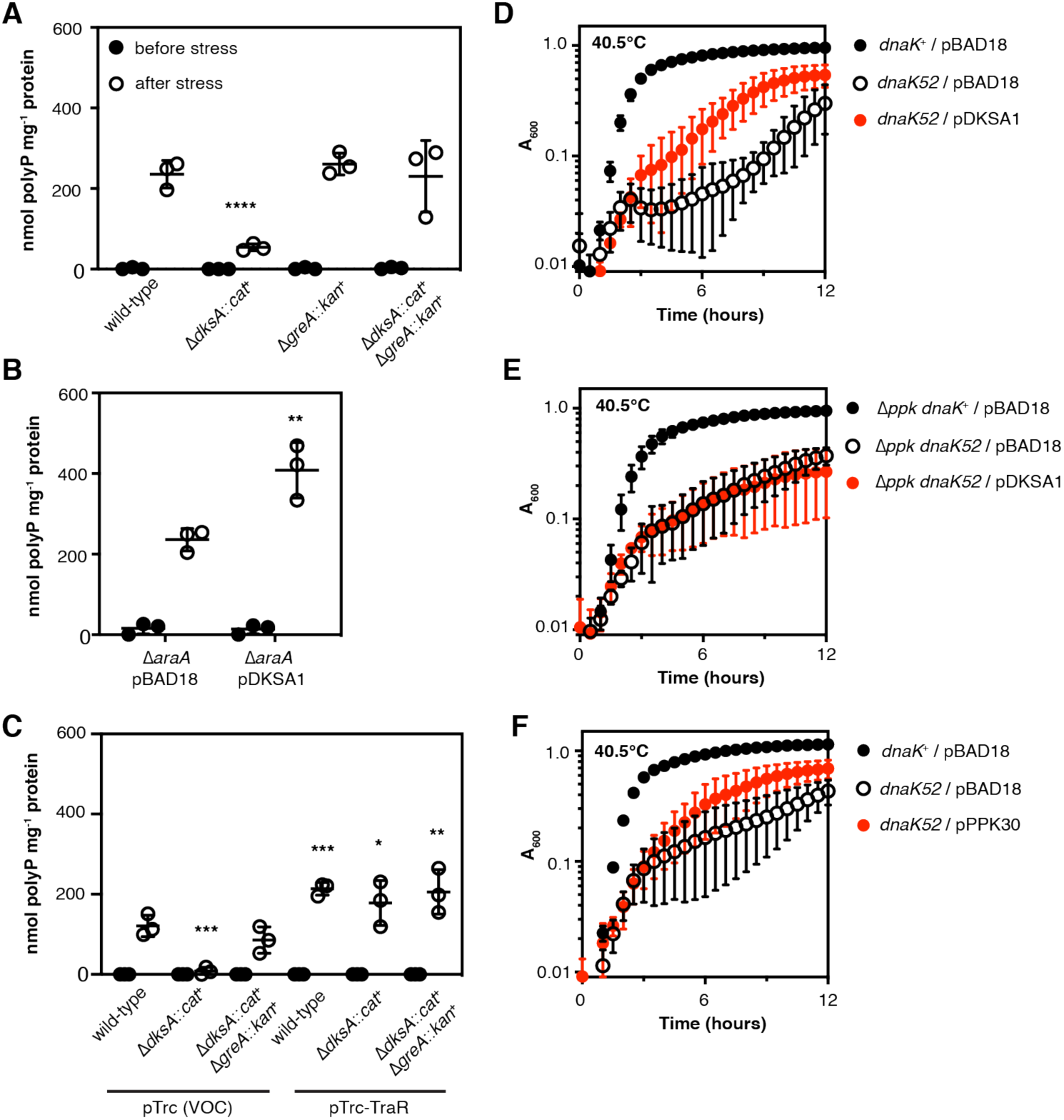
Overexpressing DksA enhances stress-induced polyP synthesis, and multicopy suppression of a *dnaK* mutation by *dksA* is polyP-dependent. (*A*) *E. coli* MG1655 wild-type and isogenic Δ*dksA1000*::*cat*^+^, Δ*greA788*::*kan*^+^, and Δ*dksA1000*::*cat*^+^ Δ*greA788*::*kan*^+^ strains were grown at 37°C to A_600_=0.2–0.4 in rich medium (LB)(black circles) and then shifted to minimal medium (MOPS with no amino acids, 4 g l^-1^ glucose, 0.1 mM K_2_HPO_4_, 0.1 mM uracil) for 2 hours (white circles)(n=3, ± SD). (*B*) MG1655 Δ*araA* containing either pBAD18 or pDKSA1 (*dksA*^+^) plasmids were grown at 37°C to A_600_=0.2–0.4 in LB containing 2 g l^-1^ arabinose (black circles) and then shifted to minimal medium containing 2 g l^-1^ arabinose for 2 hours (white circles)(n=3, ± SD). (*C*) MG1655 or isogenic Δ*dksA1000*::*cat*^+^ or Δ*dksA1000*::*cat*^+^ Δ*greA788*::*kan*^+^ mutants containing pRLG13077 (pTrc, VOC; vector-only control) or pRLG13078 (pTrc-TraR, *traR*^+^) were grown at 37°C to A_600_=0.2–0.4 in rich medium containing 1 mM IPTG (black circles) and then shifted to minimal medium containing 1 mM IPTG for 2 hours (white circles)(n=3, ± SD). PolyP concentrations are in terms of individual phosphate monomers. Asterisks indicate polyP levels significantly different from those of the wild-type control for a given experiment (two-way repeated measures ANOVA with Holm-Sidak’s multiple comparisons test, * = P<0.05, ** = P<0.01, *** = P<0.001, **** = P<0.0001). For panels (*D*) – (*E*), *E. coli* MG1655 wild-type (*dnaK*^+^) and isogenic *ΔdnaK52*::*cat*^+^, Δ*ppk-749*, or Δ*ppk-749 ΔdnaK52*::*cat*^+^ strains containing plasmids pBAD18, pDKSA1 (*dksA*^+^), or pPPK30 (*ppk*^G733A^, encoding PPK^E245K^) were grown overnight at 30°C with shaking in LB containing 100 µg ml^-1^ ampicillin, then diluted to A_600_=0.01 in LB containing 100 µg ml^-1^ ampicillin, 2 g l^-1^ arabinose, and, in (*E*), 1 mM MgCl_2_ and incubated with shaking at 40.5°C for 12 hours (n=3, ± SD).

### Multicopy suppression of a *dnaK* mutation by *dksA* is polyP-dependent

DksA is now known to be a multifunctional protein involved in genome-wide transcriptional regulation, RNA chain elongation, transcription fidelity, preventing conflicts between RNA polymerase and the DNA replisome, DNA double-strand break repair, redox sensing, and preventing RNA polymerase arrest under nutritional stress (5, 49–54), but it was originally identified as a multicopy suppressor of the heat-sensitive growth phenotype caused by a null mutation in *dnaK* (hence the name “*dksA*”: *<u>d</u>na<u>K s</u>uppressor <u>A</u>*)(55). DnaK (also known as Hsp70) is a protein-folding chaperone and central component of the *E. coli* response to proteotoxic stresses (56). Despite some speculation, the mechanism by which overexpression of DksA rescues growth of a *dnaK* mutant at high temperature has never been established (5, 52, 57). PolyP is a potent protein-stabilizing chemical chaperone (12, 58), and I wondered if the increase in polyP levels caused by DksA overexpression could account for its protective effect in a *dnaK* mutant. First, I tested the ability of a *dksA* overexpression plasmid to rescue the growth defect of a *dnaK* mutant at 40.5°C (Fig. 1D). As expected (55), it increased growth substantially, although not to wild-type levels. However, in a Δ*ppk* mutant background, in which no polyP synthesis is possible (59), overexpression of DksA provided no benefit (Fig. 1E). Expression of a hyperactive PPK^E245K^ variant (34) also improved the growth of the *dnaK* mutant (Fig. 1F), indicating that increased polyP alone is able to exert a similar protective effect as DksA overexpression. Variability was quite high in these experiments, possibly due to the rapid accumulation of suppressors reported for *dnaK* mutants (60), but these results are consistent with the idea that overproduction of polyP is at least partially responsible for *dksA*’s multicopy suppression phenotype.

### DksA requires interaction with RNA polymerase to regulate polyP synthesis

A variety of *dksA* mutant alleles have been described that encode DksA variants with clearly defined effects on DksA’s functions. I used a panel of these to determine which functions of DksA are involved in regulating polyP synthesis by complementing a *dksA* null mutant with plasmids expressing different DksA variants (Fig. 2). I tested three variants with disruptions in the conserved aspartate residues necessary for DksA’s ability to regulate transcription by destabilizing promoter open complexes (61–63). A DksA^D74E^ variant restored polyP synthesis to wild-type levels, but the more strongly defective DksA^D74N^ variant (61), expression of which is also reported to be unable to rescue growth of a Δ*dnaKJ* mutant at 42°C (57), was only able to partially restore polyP synthesis. PolyP synthesis in a strain expressing the DksA^D71N D74N^ protein, reported to be completely defective in transcriptional control (62, 63), was highly variable, but consistently lower than wild-type. These results show that the active-site aspartates are involved in but are not essential for DksA’s role in activating polyP synthesis.

**FIG 2.**
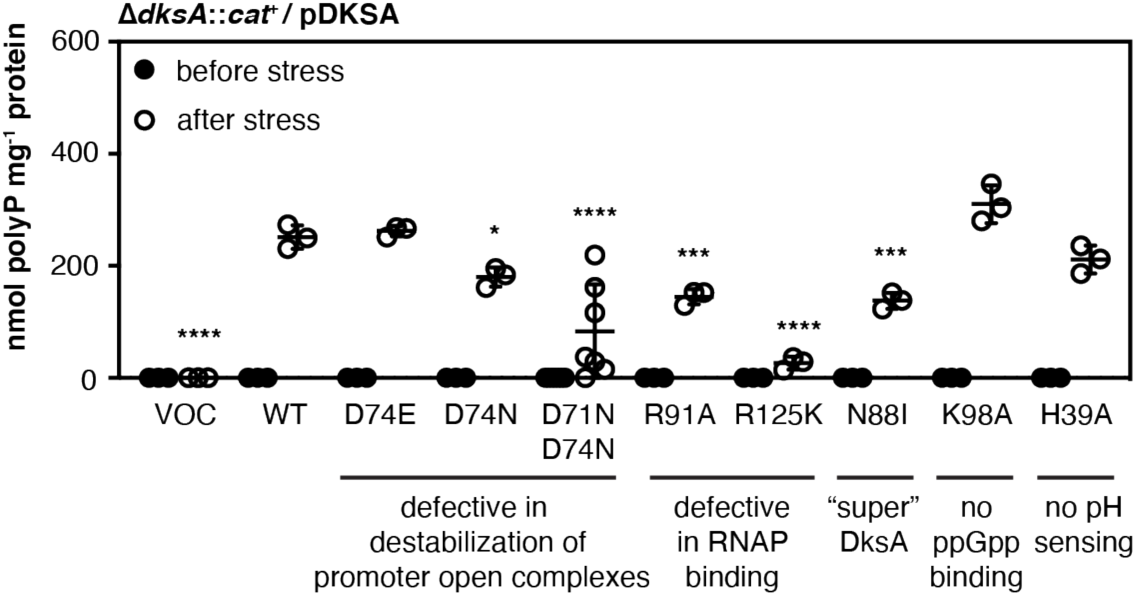
DksA requires interaction with RNA polymerase to regulate polyP synthesis. (*A*) MG1655 Δ*dksA1000*::*cat*^+^ containing pBAD18 (VOC; vector-only control), pDKSA1 (*dksA*^+^, WT), pDKSA2 (*dksA*^C222A^, D74E), pDKSA3 (*dksA*^G220A C22T^, D74N), pDKSA4 (*dksA*^C271G G272C C273G^, R91A), pDKSA5 (*dksA*^C373A G374A C375A^, R125K), pDKSA6 (*dksA*^A263T C264T^, N88I), pDKSA7 (*dksA*^A292G A293C^, K98A), pDKSA8 (*dksA*^C115G A116C C117G^, H39A), or pDKSA9 (*dksA*^G211A C213T G220A C222T^, D71N D74N) plasmids were grown at 37°C to A_600_=0.2–0.4 in rich medium (LB) containing 2 g l^-1^ arabinose and 100 µg ml^-1^ ampicillin (black circles) and then shifted to minimal medium (MOPS with no amino acids, 4 g l^-1^ glucose, 0.1 mM K_2_HPO_4_, 0.1 mM uracil) containing 2 g l^-1^ arabinose and 100 µg ml^-1^ ampicillin for 2 hours (white circles)(n=3-7, ± SD). PolyP concentrations are in terms of individual phosphate monomers. Asterisks indicate polyP levels significantly different from those of the wild-type control for each experiment (mixed effects model with Holm-Sidak’s multiple comparisons test, * = P<0.05, *** = P<0.001, **** = P<0.0001).

Mutation of the active site aspartates does not affect the binding affinity of DksA to RNA polymerase (61). Expression of a DksA^R91A^ variant, which has an approximately 2-fold reduction in binding affinity but retains some DksA functions (53, 64, 65), led to only about half the amount of polyP production seen with wild-type DksA, while expression of DksA^R125K^, which has a severe 20-fold reduction in binding affinity (65), reduced polyP accumulation 10-fold. Curiously, expression of DksA^N88I^, a so-called “super DksA” which binds to RNA polymerase about 4.5 times more tightly than wild-type DksA and has a correspondingly greater effect on expression of both positively- and negatively-regulated promoters (62) reduced polyP synthesis to a similar extent as the DksA^R91A^ variant. These results indicate that DksA must bind to RNA polymerase to activate polyP synthesis, but that either increases or decreases in its binding affinity can reduce the extent of that synthesis. This generally supports the hypothesis that DksA may influence polyP synthesis by controlling the access of other regulators (*e.g.* GreA) to the secondary channel of RNA polymerase (11, 38).

As expected, since a *relA spoT* mutant incapable of making ppGpp is able to synthesize wild-type levels of polyP under these conditions (11), expression of DksA^K98A^, which cannot bind ppGpp (64), allowed wild-type levels of polyP synthesis. DksA activity is sensitive to cytoplasmic pH (66), but disruption of the pH-sensing histidine residue (DksA^H39A^) also had no inhibitory effect on polyP synthesis. DksA contains a redox-sensing zinc finger domain (52, 67), but I was unable to test whether this plays a role in polyP synthesis in this set of experiments, since mutations disrupting this domain destabilize the protein and are equivalent to Δ*dksA* mutations (68).

### Levels of *ppk* and *ppx* mRNA decrease after nutrient limitation stress

I have previously used reporter fusions to show that expression from the promoter of the *ppk-ppx* operon (P*ppk*) is decreased in the absence of either DksA or ppGpp, but does not change significantly after nutrient limitation stress in wild-type, *dksA*, or *dksA greA* mutant strains (11). However, since both DksA and GreA can act as transcription elongation factors (50, 51, 69, 70), I hypothesized that the effect of *dksA* and *greA* mutations on polyP accumulation might result from changes in *ppk* or *ppx* transcript accumulation independent of transcription initiation from P*ppk*. I therefore used quantitative reverse transcription PCR (qRT-PCR) to measure the amount of mRNA from three different points in the *ppk*-*ppx* operon (Fig. 3A) before and after nutrient limitation stress in wild-type, *dksA*, and *dksA greA* strains.

**FIG 3.**
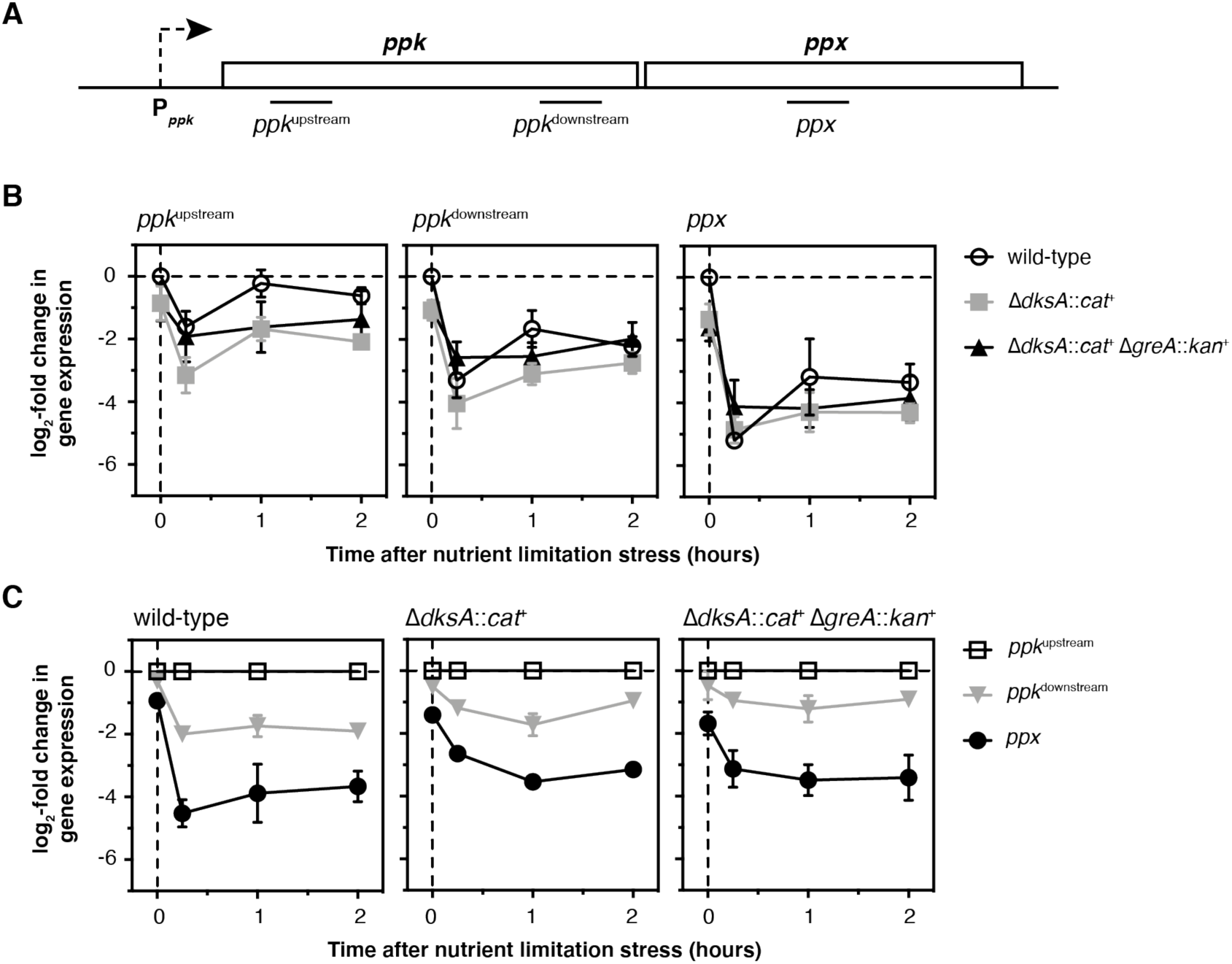
Levels of *ppk* and *ppx* mRNA decrease after nutrient limitation stress. (*A*) Position of qPCR amplicons within the *ppk* and *ppx* genes. (*B, C*) *E. coli* MG1655 wild-type and isogenic Δ*dksA1000*::*cat*^+^, and Δ*dksA1000*::*cat*^+^ Δ*greA788*::*kan*^+^ strains were grown at 37°C to A_600_=0.2–0.4 in rich medium (LB) and then shifted to minimal medium (MOPS with no amino acids, 4 g l^-1^ glucose, 0.1 mM K_2_HPO_4_, 0.1 mM uracil) for 2 hours. qRT-PCR was used to measure fold changes in transcript abundance at the indicated timepoints (n=3, ± SD). In (*B*), changes in expression for each amplicon are normalized to expression of that amplicon in the wild-type strain before stress treatment (t = 0 h). In (*C*), changes in expression for each amplicon are normalized to expression of the *ppk*^upstream^ amplicon in the same strain at each time point.

As I reported previously using promoter fusions (11), expression of *ppk* in the absence of stress was about 2-fold lower in the *dksA* and *dksA greA* mutants than in the wild-type (Fig. 3B). Surprisingly, however, I found that in all three strains, levels of transcripts from both the 5’ and 3’ ends of the *ppk* gene, referred to hereafter as *ppk*^upstream^ (nucleotides 209-409 of *ppk*) and *ppk*^downstream^ (nucleotides 1530-1725 of *ppk*), rapidly decreased 4 to 16-fold after stress, and levels of *ppx* transcripts (nucleotides 510-695 of *ppx*) decreased more than 32-fold. Normalizing the levels of *ppk*^downstream^ and *ppx* transcripts to the amount of *ppk*^upstream^ transcripts at each time point (Fig. 3C) showed that in the absence of stress, both ends of the *ppk* transcript are present at the same level, with 2 to 4-fold fewer *ppx* transcripts present. After stress, however, levels of the *ppk*^downstream^ transcript drop 2 to 4-fold below the levels of the *ppk*^upstream^ transcript, while amounts of *ppx* transcript decrease more dramatically (8 to 32-fold). These results indicate that nutrient limitation stress has a significant effect on the accumulation of *ppk* and *ppx* transcripts and on their relative levels in the cell, and demonstrate previously unsuspected regulation of transcript elongation or mRNA stability in the *ppk-ppx* operon. However, since there were no dramatic differences in these patterns between wild-type, *dksA*, and *dksA greA* strains, and certainly no effects that correlate with the respective polyP accumulation in those strains, these results do not account for how DksA and GreA regulate polyP accumulation during nutrient limitation.

In parallel with the above experiments, I also measured levels of 16S ribosomal RNA (*rrsD*) at each time point after nutrient limitation (Supplemental Figure S1). Amino acid starvation is a well-known inducer of the stringent response (71), in which DksA and ppGpp repress rRNA transcription (5), but the specific nutrient limitation stress used here (shift of exponentially-growing cultures from LB to MOPS minimal medium containing 4 g l^-1^ glucose and 0.1 mM K_2_HPO_4_), originally described by Arthur Kornberg’s lab as a stress condition that robustly stimulates polyP synthesis (10), actually led to a 1.5 to 3-fold increase in *rrsD* expression. This is consistent with the lack of a role for ppGpp in polyP regulation under these conditions (11), but does reopen the question of what roles ppGpp and DksA might play in polyP accumulation under conditions that do activate the classic stringent response.

### Identifying genes regulated by DksA and GreA under nutrient limitation conditions

I next used mRNA sequencing of wild-type, *dksA*, and *dksA greA* strains immediately before and 5 minutes after nutrient starvation stress to screen for genes whose expression patterns correlated, either positively or negatively, with the accumulation of polyP by these strains (Supplemental Dataset S1, Fig. S2). Both DksA and GreA are global regulators (38), and nutrient limitation induced sweeping changes in gene expression in all three strains, so I found there to be a surprisingly large number of such genes. I used qRT-PCR to validate several of the most striking candidates (Fig. 4A-C). This reproduced the results of the RNA sequencing screen in all cases, and confirmed that expression of genes involved in the regulation and synthesis of flagella (72, 73)(Supplemental Fig. S3A) was inversely correlated with capacity for polyP synthesis, both before and after stress (Fig. 4A). Conversely, the expression of most genes involved in glycerol dissimilation (72, 74)(Supplemental Figs. S3B,C) was positively correlated with capacity for polyP synthesis, at least before stress (Fig. 4B,C). Finally, the expression of the *gadE* operon, which encodes a central regulator of the *E. coli* acid stress response (72, 75) and the MdtEF multidrug exporter (76)(Supplemental Figure S3D) was inversely correlated with capacity for polyP synthesis both before and after stress. However, no mutations blocking the expression of operons encoding genes for flagella (Fig. 4D), glycerol dissimilation (Fig. 4E, F), or acid stress response (Fig. 4F) had any effect on polyP synthesis.

**FIG 4.**
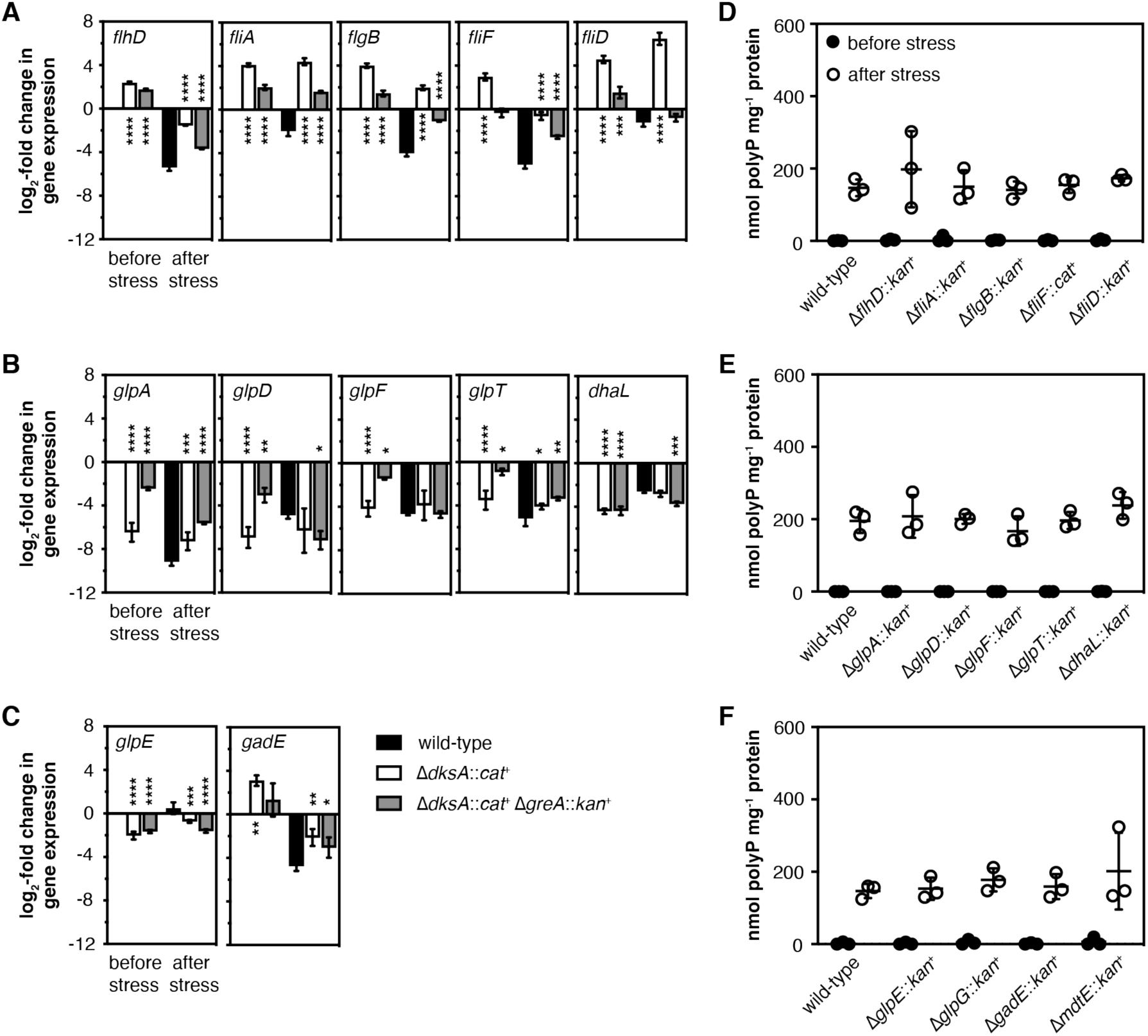
Mutations disrupting *dksA*- and *greA*-regulated operons do not affect polyP accumulation. *E. coli* MG1655 wild-type and isogenic (*A-C*) Δ*dksA1000*::*cat*^+^, and Δ*dksA1000*::*cat*^+^ Δ*greA788*::*kan*^+^, (*D*) Δ*flhD745*::*kan*^+^, Δ*fliA*::*kan*^+^, Δ*flgB742*::kan^+^, Δ*fliF1000*::*cat*^+^, and Δ*fliD770*::*kan*^+^, (*E*) Δ*glpA721*::*kan*^+^, Δ*glpD759*::*kan*^+^, Δ*glpF786*::*kan*^+^, Δ*glpT720*::*kan*^+^, and Δ*dhaL789*::*kan*^+^, or (*F*) Δ*glpE758*::*kan*^+^, Δ*glpG757*::*kan*^+^, Δ*gadE767*::*kan*^+^, and Δ*mdtE768*::*kan*^+^ strains were grown at 37°C to A_600_=0.2–0.4 in rich medium (LB)(black circles) and then shifted to minimal medium (MOPS with no amino acids, 4 g l^-1^ glucose, 0.1 mM K_2_HPO_4_) for 2 hours (white circles)(n=3, ± SD). (*A-C*) qRT-PCR was used to measure fold changes in transcript abundance of the indicated genes before and 15 min after nutrient limitation, relative to expression in the wild-type strain in the absence of stress (n=3, ± SD). (*D-F*) PolyP concentrations are in terms of individual phosphate monomers. Asterisks indicate gene expression significantly different from those of the wild-type control at each time point (two-way repeated measures ANOVA with Holm-Sidak’s multiple comparisons test, * = P<0.05, ** =P<0.01, *** = P<0.001, **** = P<0.0001). No significant differences were found between polyP levels of any strain and the wild-type control.

### Does overexpression of flagellar regulators inhibit polyP synthesis?

Although knocking out flagellar genes had no effect on polyP synthesis (Fig. 4D), this did not necessarily prove that the inhibition of polyP synthesis in a *dksA* mutant was not due to overexpression of one of the flagellar regulators: FlhDC, FliA, or FliZ (73, 77–79). Expression of FlhDC responds to a wide variety of environmental stresses (79, 80) and activates expression of flagellar genes (including *fliA* and *fliZ*)(Supplemental Fig. S3A), as well as a number of genes with other functions (73). Similarly, FliA (also known as *σ*^28^, *σ*^F^, or RpoF) is a sigma factor that activates expression of many genes, not all of which are involved in motility (73). FliZ is a transcription factor, cotranscribed with FliA, that competes for promoter binding with RpoS, the master regulator of the general stress response in *E. coli* (4, 81), so could directly or indirectly affect the expression of hundreds of genes (77, 78). A *dksA* mutant was hypermotile, as expected (82), and a *dksA greA* double mutant was almost completely non-motile (Fig. 5A). Stringent mutations of RNA polymerase (83, 84) that lead, by an unknown mechanism presumably related to that of a *dksA* mutant, to reduced polyP accumulation (11) also had increased motility (Supplemental Fig. S4), despite growing substantially more slowly than the wild-type (85). However, overexpression of FlhDC, FliA, or FliZ did not repress polyP synthesis (Fig. 5B), nor was the inhibitory effect of a *dksA* mutation suppressed by *flhDC* or *fliAZ* mutations (Fig. 5C). Repression of polyP synthesis in a *dksA* mutant is therefore not due, directly or indirectly, to overproduction of any of the flagellar regulators. Surprisingly, though, mutation of either *flhD* or *fliA* did prevent overexpression of DksA from enhancing polyP synthesis (Fig. 5D), suggesting that the flagellar regulators might have an indirect effect on polyP regulation by DksA. This is not unprecedented, since work from several laboratories has shown indirect regulation of promoters driven by alternative sigma factors by both ppGpp and DksA, which is thought to depend on changes in the availability of RNA polymerase core enzyme due to DksA/ppGpp-dependent repression of rRNA transcription (86–91). FliA is abundant and has high affinity for RNA polymerase (92–94), so deletion of *fliA* could plausibly have a substantial effect on the equilibrium among other sigma factors and core RNA polymerase (95, 96).

**FIG 5.**
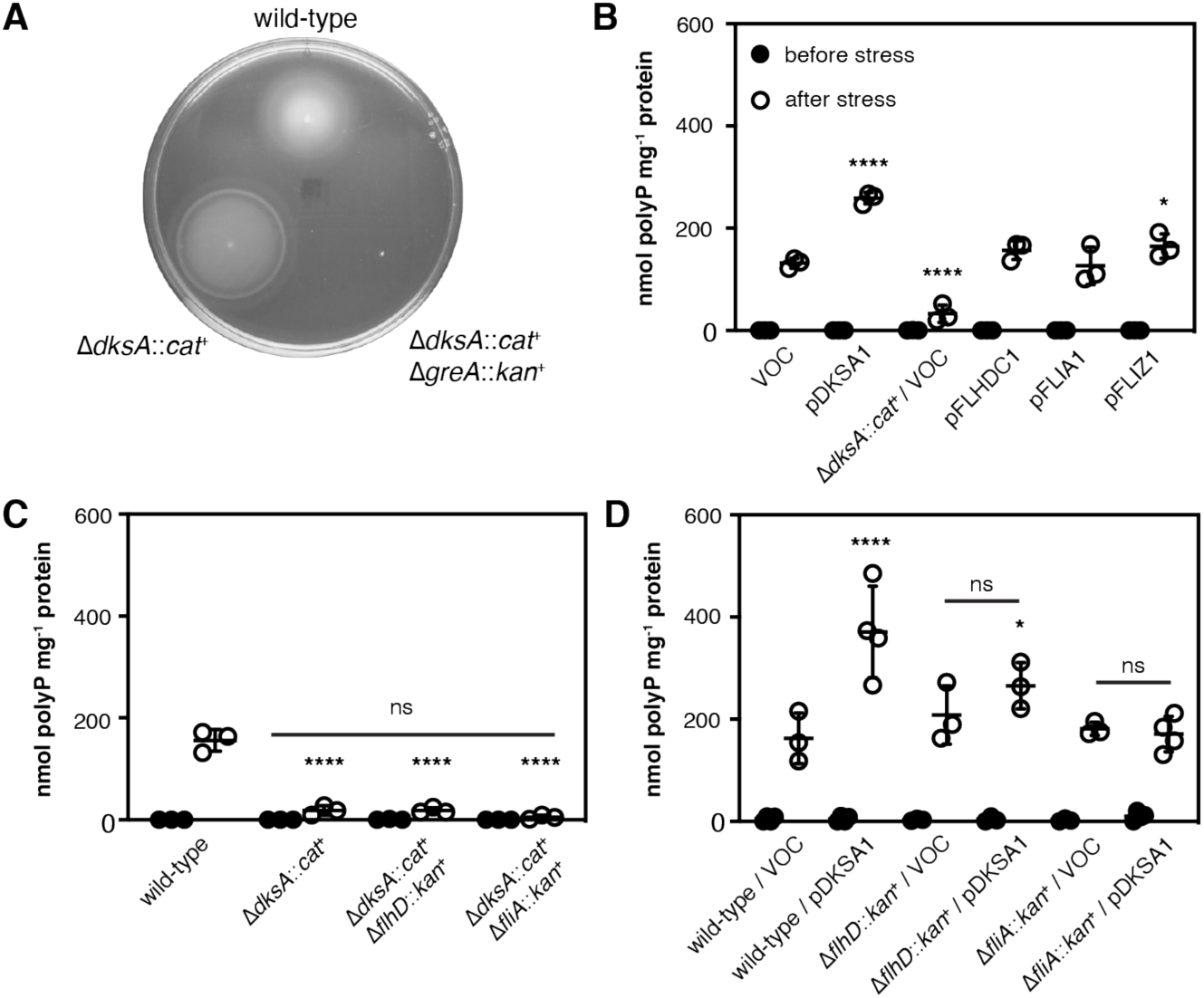
Suppression of polyP synthesis in a *dksA* mutant is not due to overexpression of flagellar regulators. (*A*) *E. coli* MG1655 wild-type and isogenic Δ*dksA1000*::*cat*^+^ and Δ*dksA1000*::*cat*^+^ Δ*greA788*::*kan*^+^ strains were inoculated into LB containing 0.25% agar and incubated at room temperature (representative of 3 independent experiments). (*B*) – (*D*) *E. coli* MG1655 wild-type or isogenic Δ*dksA1000*::*cat*^+^, Δ*dksA1000*::*cat*^+^ Δ*flhD745*::*kan*^+^, Δ*dksA1000*::*cat*^+^ Δ*fliA*::*kan*^+^, Δ*flhD745*::*kan*^+^, or Δ*fliA*::*kan*^+^ strains containing, where indicated, pBAD18 (VOC; vector-only control) or the indicated pBAD18-derived plasmids were grown at 37°C to A_600_=0.2–0.4 in rich medium (LB) containing, when plasmids were present, 100 µg ml^-1^ ampicillin (black circles) and then shifted to minimal medium (MOPS with no amino acids, 4 g l^-1^ glucose, 0.1 mM K_2_HPO_4_, 0.1 mM uracil) containing 100 µg ml^-1^ ampicillin for 2 hours (white circles)(n=3-4, ± SD). Experiments in panels (*B*) and (*D*) included 2 g l^-1^ arabinose in both rich and minimal media. PolyP concentrations are in terms of individual phosphate monomers. Asterisks indicate polyP levels significantly different from those of the wild-type control for each experiment (two-way repeated measures ANOVA or mixed-effects model with Holm-Sidak’s multiple comparisons test, * = P<0.05, **** = P<0.0001). No significant differences (ns) were found among the three indicated strains in (*C*) or between the indicated VOC and pDKSA1-containing strains in (*D*).

### The role of alternative sigma factors in regulation of polyP accumulation

In addition to FliA and RpoS, *E. coli* has three other alternative sigma factors that also respond to environmental stresses and drive the expression of large numbers of genes: RpoH (*σ*^32^, *σ*^H^), which controls chaperones and proteases in response to protein unfolding stresses (81, 97), RpoN (*σ*^54^, *σ*^N^), which responds to a number of signals, including nitrogen starvation (98), and RpoE (*σ*^24^, *σ*^E^), which responds to perturbations in the cell envelope (99). The only other sigma factor in *E. coli*, FecI (*σ*^19^), responds to ferric citrate and controls expression of only a single promoter (that of *fecA*)(100). Regulation of gene expression in response to changes in environmental conditions involves complex, overlapping interactions among these sigma factors and other stress’response pathways, including the stringent response (4, 77, 81, 82, 86–88, 90, 101). The observation that overexpression of the RpoS antagonist FliZ slightly increased polyP accumulation (Fig. 5B) and that a *dksA fliA*::*kan*^+^ mutant, which lacks both *fliA* and *fliZ*, appeared to contain even less polyP than a *dksA* mutant (Fig. 5C)(although that difference was not statistically significant in this set of experiments), supported the idea that alternative sigma factors might be playing a role in modulating polyP synthesis. While I have not found RpoS to be required for polyP accumulation in response to the nutrient limitation stress used here (11), others have reported roles for either RpoS or RpoN under some other stress conditions (10). Regardless, I wanted to know whether any of *E. coli*’s other major stress response pathways were interacting with DksA in the control of polyP synthesis. I therefore performed a series of epistasis experiments to determine the role of each of the above alternative sigma factors in polyP accumulation by *E. coli* in the presence and absence of DksA (Fig. 6).

**FIG 6.**
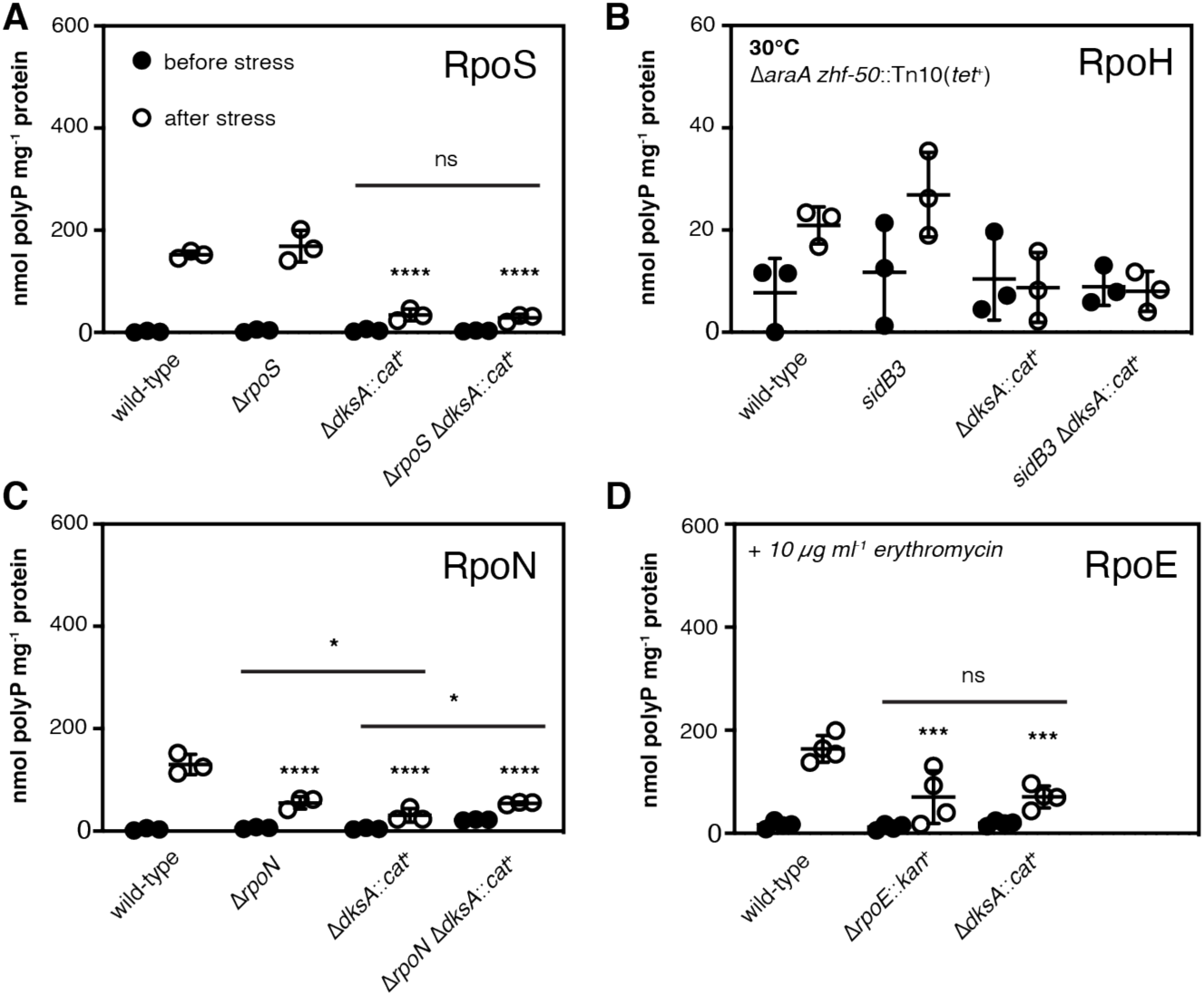
Roles of stress-sensing alternative sigma factors in polyP synthesis. *E. coli* MG1655 wild-type and isogenic Δ*rpoS746,* Δ*dksA1000*::*cat^+^,* Δ*rpoS746* Δ*dksA1000*::*cat^+^*, Δ*araA zhf-50*::Tn10(*tet*^+^), Δ*araA zhf-50*::Tn10(*tet*^+^) *sidB3,* Δ*araA zhf-50*::Tn10(*tet*^+^) Δ*dksA1000*::*cat^+^,* Δ*araA zhf-50*::Tn10(*tet*^+^) *sidB3* Δ*dksA1000*::*cat^+^,* Δ*rpoN730,* Δ*rpoN730* Δ*dksA1000*::*cat^+^,* or Δ*rpoE1000*::*kan*^+^ strains were grown at 37°C or 30°C, as indicated, to A_600_=0.2–0.4 in rich medium (LB) containing 100 µg ml^-1^ ampicillin (black circles) and then shifted to minimal medium (MOPS with no amino acids, 4 g l^-1^ glucose, 0.1 mM K_2_HPO_4_, 0.1 mM uracil) containing 100 µg ml^-1^ ampicillin for 2 hours (white circles)(n=3-4, ± SD). Experiments in panel (*D*) included 10 µg ml^-1^ erythromycin in both rich and minimal media. PolyP concentrations are in terms of individual phosphate monomers. Unless specifically indicated, asterisks indicate polyP levels significantly different from those of the wild-type control for each experiment (two-way repeated measures ANOVA with Holm-Sidak’s multiple comparisons test, ns = P>0.05, * = P<0.05, *** = P<0.001, **** = P<0.0001).

As I have previously reported (11), an *rpoS* mutant had no defect in polyP production under these conditions, and polyP levels in an *rpoS dksA* mutant were indistinguishable from those in a *dksA* mutant (Fig. 6A). *E. coli m*utants lacking *rpoH* are not viable above 20°C, although they rapidly accumulate suppressors that allow them to grow at 30°C or higher (102, 103). To assess the possible role of RpoH in polyP synthesis, I therefore used the defective *rpoH* allele *sidB3*, which contains much less RpoH than wild-type and cannot induce the expression of heat shock proteins in response to elevated temperature, but grows stably at 30°C (60). PolyP production in all strains (including MG1655 wild-type; data not shown) was roughly 10-fold lower at 30°C than at 37°C, indicating that temperature strongly influenced polyP production and making comparisons between strains difficult, but the pattern of polyP production in *sidB3* and *sidB3 dksA* double mutants did not suggest that *sidB3* has a major effect on the *dksA* phenotype (Fig. 6B). Mutants lacking *rpoN* had a significant defect in polyP synthesis (Fig. 6C), as did an *rpoN dksA* double mutant. RpoE is essential in *E. coli* MG1655 under normal growth conditions, but can be stably knocked out in the presence of subinhibitory concentrations of erythromycin (104, 105). Under these conditions, both the *dksA* and the *rpoE* mutant had significant polyP synthesis defects (Fig. 6D). Despite multiple attempts, I was unable to construct an *rpoE dksA* double mutant, indicating that these genes are synthetically lethal. Deletion of *rsd*, encoding a negative regulator of RpoD that facilitates sigma factor competition during stationary phase (86, 106) had no effect on polyP synthesis (Supplemental Figure S5).

Overexpressing DksA was able to restore polyP synthesis to the *rpoN* mutant, but not to the *rpoE* mutant (Fig. 7A). Conversely, overexpression of RpoE increased polyP production in the wild-type and the *rpoN* mutant, but not in the *dksA* or, surprisingly, the *rpoE* mutant (Fig. 7B). The meaning of this last result is unclear, but the Δ*rpoE*::*kan*^+^ mutant did not tolerate overexpression of *rpoE* very well and had a very long lag phase under even modest induction conditions (data not shown), possibly because of the lack of the RseA anti-sigma factor in this strain (107). However, since *rpoE* mutants are inviable at 42°C (108), I was unable to resolve the Δ*rpoE*::*kan*^+^ insertion (109) to test this idea. Nevertheless, the results of this experiment indicate that both *rpoE* and *dksA* are positive regulators of polyP synthesis upon nutrient limitation and depend on one another to have this effect, but that overexpression of either gene rescues polyP synthesis in an *rpoN* mutant.

**FIG 7.**
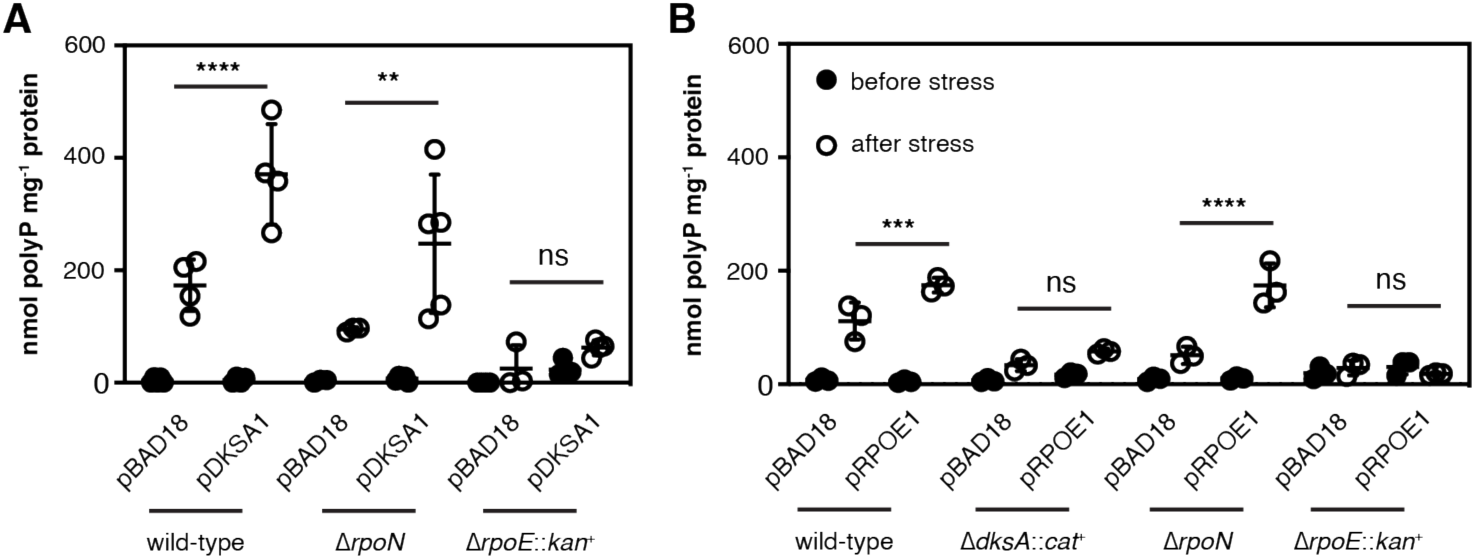
Complementation of polyP-deficient mutants with ectopically-expressed DksA and RpoE. *E. coli* MG1655 wild-type and isogenic Δ*dksA1000*::*cat^+^,* Δ*rpoN730,* or Δ*rpoE1000*::*kan*^+^ strains containing pBAD18 or the indicated pBAD18-derived plasmids were grown at 37°C to A_600_=0.2–0.4 in rich medium (LB) containing 100 µg ml^-1^ ampicillin (black circles) and 0.2% (*A*) or 0.0125% (*B*) arabinose and then shifted to minimal medium (MOPS with no amino acids, 4 g l^-1^ glucose, 0.1 mM K_2_HPO_4_, 0.1 mM uracil) containing 100 µg ml^-1^ ampicillin and 0.2% (*A*) or 0.0125% (*B*) arabinose for 2 hours (white circles)(n=3-5, ± SD). Growth media for *rpoE* mutants included 10 µg ml^-1^ erythromycin in both rich and minimal media. PolyP concentrations are in terms of individual phosphate monomers. Unless specifically indicated, asterisks indicate polyP levels significantly different from those of the wild-type control for each experiment (two-way repeated measures ANOVA or mixed-effects model with Holm-Sidak’s multiple comparisons test, ns = P>0.05, ** = P<0.01, *** = P<0.001, **** = P<0.0001).

## DISCUSSION

The purpose of this work was to characterize the role of DksA in regulating polyP synthesis in *E. coli*, with the ultimate goal of understanding how exposure to stressful environmental conditions leads to polyP accumulation. While I have made some progress towards this goal, the experiments presented here have also revealed several layers of unexpected complexity and emphasize the fact that polyP regulation is deeply entwined in the complicated and multifactorial general stress response of *E. coli*.

While changes in DksA expression did not affect polyP levels under non-stress conditions, the amount of polyP produced after nutrient limitation correlated well with the amount of DksA being expressed (Fig. 1A,B). This also held true for expression of TraR, which has many of the same effects on RNA polymerase as DksA and ppGpp (47)(Fig. 1C). Experiments with *dnaK* knockouts indicated that the original multicopy suppression phenotype that gave DksA its name (55) is, in fact, probably due to the polyP overproduction that results from DksA overexpression (Fig. 1D-F). PolyP overproduction could potentially also play a role in explaining other phenotypes associated with overexpression of *dksA*, including multicopy suppression of *grpE*, *prc*, *yhhP*, and *mukB* mutations (55, 110–112), although since DksA has many documented functions (5, 49–54), these would need to be tested on an individual basis. The ability of DksA to induce polyP accumulation was influenced its ability to bind RNA polymerase, but was largely independent of other functional residues (Fig. 2), which may help to explain the suppression of the *dksA* mutant phenotype by a *greA* deletion (Fig. 1A). Both DksA and GreA bind in the secondary channel of RNA polymerase, and changes in the equilibrium between these two proteins at that site are thought to have regulatory consequences at certain promoters, although the details have not yet been fully worked out (35, 37, 38, 40, 41).

The nutrient shift stress used in this and earlier works (11, 13, 30, 34) was originally described by Arthur Kornberg’s lab (10) and is a robust and reproducible way to induce substantial polyP synthesis in *E. coli*. Despite its technical simplicity, however, the results of the experiments in this paper show that “nutrient limitation” is physiologically complex and leads to dramatic, genome-wide changes in gene expression, many of which, like polyP production, are strongly affected by *dksA* and *greA* mutations (Supplemental Dataset S1, Fig. S2). These include up- or down-regulation of operons regulated by many of the known stress-responsive regulators in *E. coli*, including multiple alternative sigma factors and transcription factors, and teasing apart which of these regulons directly influence polyP production is a significant challenge (Fig. 4,5). The assumption that this nutrient shift induces a stringent response (13) also appears to be wrong, since rRNA expression does not decrease after the shift (5)(Supplemental Fig. S1). Typically, the ppGpp-dependent stringent response to amino acid starvation is induced in laboratory conditions by addition of serine hydroxamate or specific isoleucine limitation (5, 113, 114), which is different from the simultaneous complete amino acid deprivation and phosphate limitation used here (10). As mentioned above, this may explain why ppGpp is not required for polyP synthesis under these conditions (11), but reopens the question of what, if any, role ppGpp plays in polyP regulation under other stress conditions. PolyP accumulation can reportedly be induced in *E. coli* by various kinds of starvation, serine hydroxamate, phosphate starvation or surplus, entry into stationary phase, hypochlorous acid, salt, or heat (10, 12, 58, 115). A signal common to all of these conditions is not obvious, but my results with alternative sigma factor mutants indicate that multiple stress-sensing mechanisms feed into control of polyP, either directly or indirectly.

*E. coli* encodes seven sigma factors: the housekeeping sigma RpoD and the alternative sigma factors RpoS, RpoN, RpoE, RpoH, FliA, and FecI (93, 116). Each alternative sigma factor drives expression of genes in response to particular environmental stressors (*e.g.* RpoS is active during stationary phase and RpoH is active during heat shock)(4, 81, 97), but there are complex interactions among their regulons that are not fully understood. Expression of *rpoN* is activated by RpoE (117), expression of *rpoE* can be activated by RpoN or RpoS (118), both RpoN and RpoE activate transcription of *rpoH* (119, 120), and the FliA-dependent regulator FliZ inhibits expression of RpoS-dependent promoters (77, 78). DksA and ppGpp can influence many of these interactions, as well. Expression of *fliA* is inhibited by DksA (82), and DksA and ppGpp are required for transcription of at least some RpoN and RpoE-dependent promoters (87, 101). Some of these effects are indirect and appear to depend on competition of sigma factors for core RNA polymerase (95, 96). For example, deletion of *rpoN* enhances expression from RpoS-dependent promoters, and the enhanced motility of an *rpoS* mutant is both FliA- and RpoN-dependent (88). Expression from weak (but not strong) RpoN-dependent promoters requires DksA and ppGpp *in vivo*, but not *in vitro* (87), but ppGpp facilitates RpoS and RpoH competition with RpoD for RNA polymerase both *in vitro* and *in vivo* (86). No simple model can currently explain all of these interactions, which appear to be both promoter- and growth condition-specific (101).

In light of this complexity, my results showing the impacts of DksA, RpoN, and RpoE on polyP production (Figs. 1, 6, 7) must be interpreted cautiously. The effect of a *fliA* knockout on the DksA overexpression phenotype (Fig. 5D) hints at a role for sigma factor competition in that mutant background, but whether this is physiologically important in a wild-type strain is unclear. Complementation analysis (Fig. 7) shows that increasing expression of either RpoE or DksA can compensate for the loss of RpoN, but that both RpoE and DksA are necessary for polyP accumulation. RpoN-dependent promoters are not thought to directly require DksA (87), but some RpoE-dependent promoters do (101). On the other hand, RpoN-dependent promoters do require enhancer proteins, of which *E. coli* has 11 (98). A mutant lacking the GlnG (NtrC) enhancer protein, reported to be required for polyP production upon nitrogen exhaustion (10), has no defect in polyP accumulation after nutrient limitation (11), but the others have not been tested yet. Future experiments will be needed to explore how the well-characterized inputs to the RpoN and RpoE regulons (98, 99, 121) contribute to polyP accumulation. Perhaps the simplest model to explain the current data is that there is a signaling cascade leading to polyP synthesis in which an RpoN-dependent gene acts upstream of an RpoE/DksA-dependent gene, but without knowing which regulated genes directly affect polyP synthesis, it is difficult to make definitive conclusions about the nature of the regulatory network involved. Identifying these gene(s) is a major focus of my lab’s future directions.

It is clear from this and previous work that transcription of *ppk* itself does not increase upon polyP-stimulating stresses in *E. coli* (11, 34). In fact, while transcription initiation from the P*ppk* promoter stays fairly constant before and after stress (11, 34), I have now found that the levels of *ppk* and *ppx* mRNA transcripts drop precipitously after nutrient limitation (Fig. 3). The mechanism underlying this drop is currently unknown. It is reduced slightly by deletion of *dksA*, but the *dksA* and *dksA greA* mutants exhibit similar amounts of expression, so the regulation of *ppk-ppx* transcript elongation or mRNA stability cannot account for the differences in polyP accumulation in those mutants (Fig. 1A). Deletion of *ppx* does not eliminate stress regulation of polyP production (11, 12, 34). There must, therefore, be post-transcriptional regulation of PPK activity to account for increased polyP production. This, presumably, is what is being controlled by RpoN, RpoE, and DksA. The existence of *ppk*\* mutations which change amino acids distant from the active site of PPK and result in massive overaccumulation of polyP *in vivo* (like PPK^E245K^; Fig. 1F)(34) suggests that the PPK enzyme itself may be controlled by post-translational or allosteric mechanisms, and work is ongoing in my lab to determine how PPK activity is modulated.

The idea that there must be an unknown factor or factors regulating PPK activity is not new. Kornberg hypothesized the existence of such a factor, which he called “X”, in 1998, to explain the varying requirements for different regulators he observed under different stress conditions (10). My experiments have now added several new pieces of information to this old puzzle. The involvement of multiple stress sensing systems helps to explain how so many different stress conditions can lead to polyP production and why different regulators may be required to respond to different stressors, but the focus now must be on identifying the mysterious factor “X” and understanding how it ties polyP regulation into the rest of the *E. coli* stress response machinery.

## MATERIALS AND METHODS

### Bacterial strains, growth conditions, and molecular methods

All strains and plasmids used in this study are listed in Table 1. I carried out DNA manipulations by standard methods (122, 123) in the *E. coli* cloning strain DH5*α* (Invitrogen). I grew *E. coli* at the indicated temperatures in Lysogeny Broth (LB)(124) containing 5 g l^-1^ NaCl and, where indicated, 1 mM MgCl_2_ (34). I added antibiotics when appropriate: ampicillin (100 µg ml^-1^), chloramphenicol (17 or 34 µg ml^-1^), erythromycin (10 µg ml^-1^), kanamycin (25 or 50 µg ml^-1^), rifampicin (50 µg ml^-1^), spectinomycin (50 µg ml^-1^), or tetracycline (15 µg ml^-1^).

**TABLE 1.**
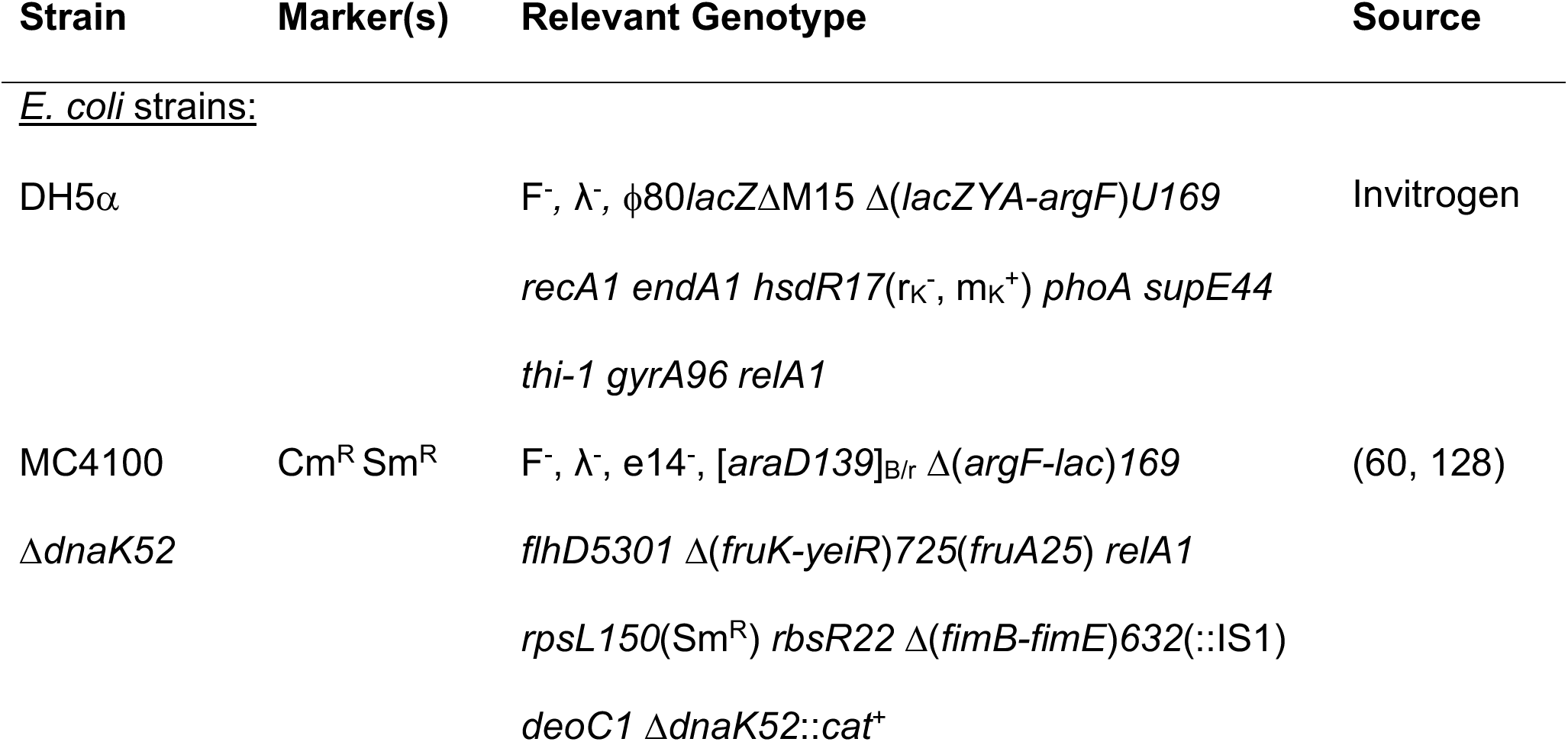
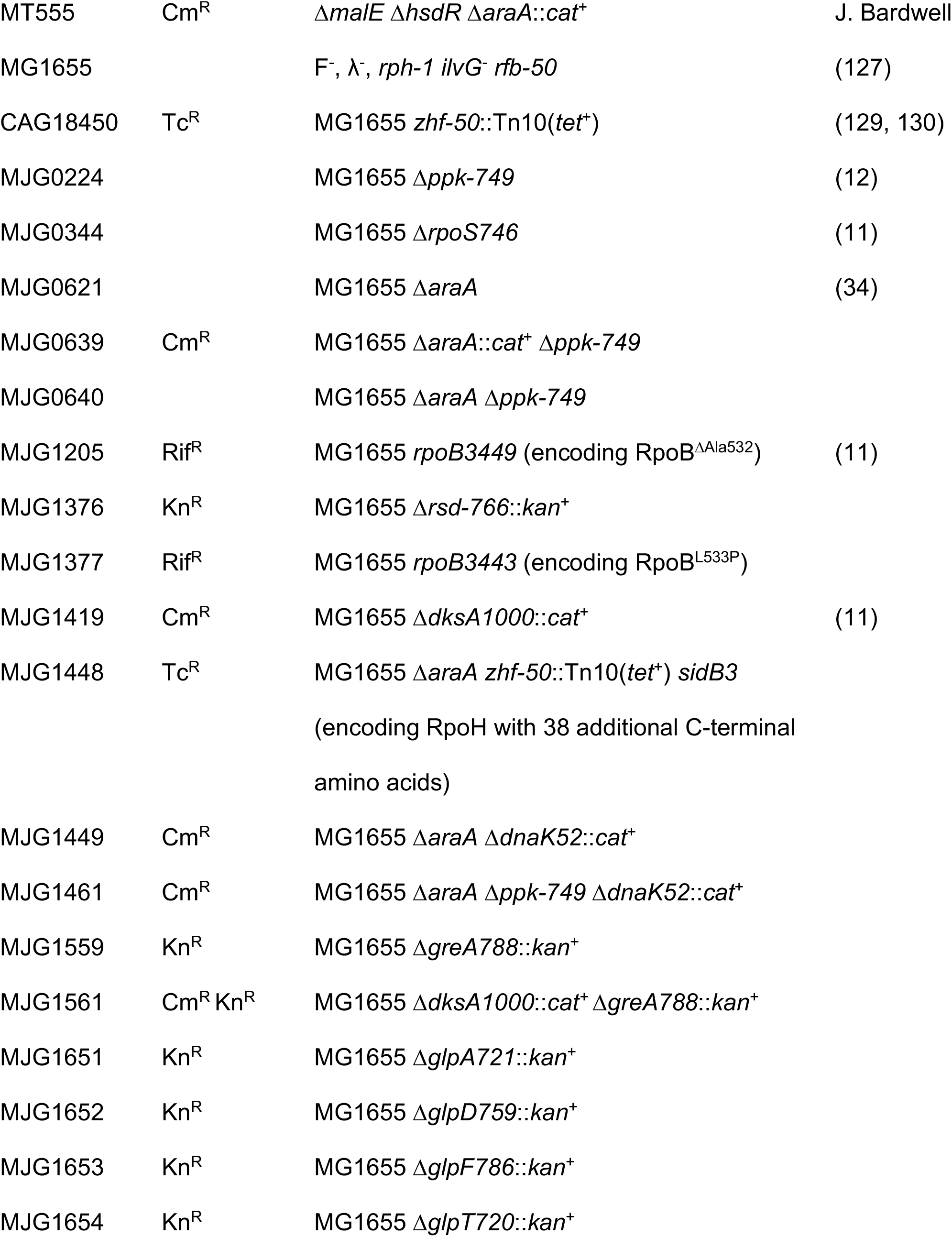
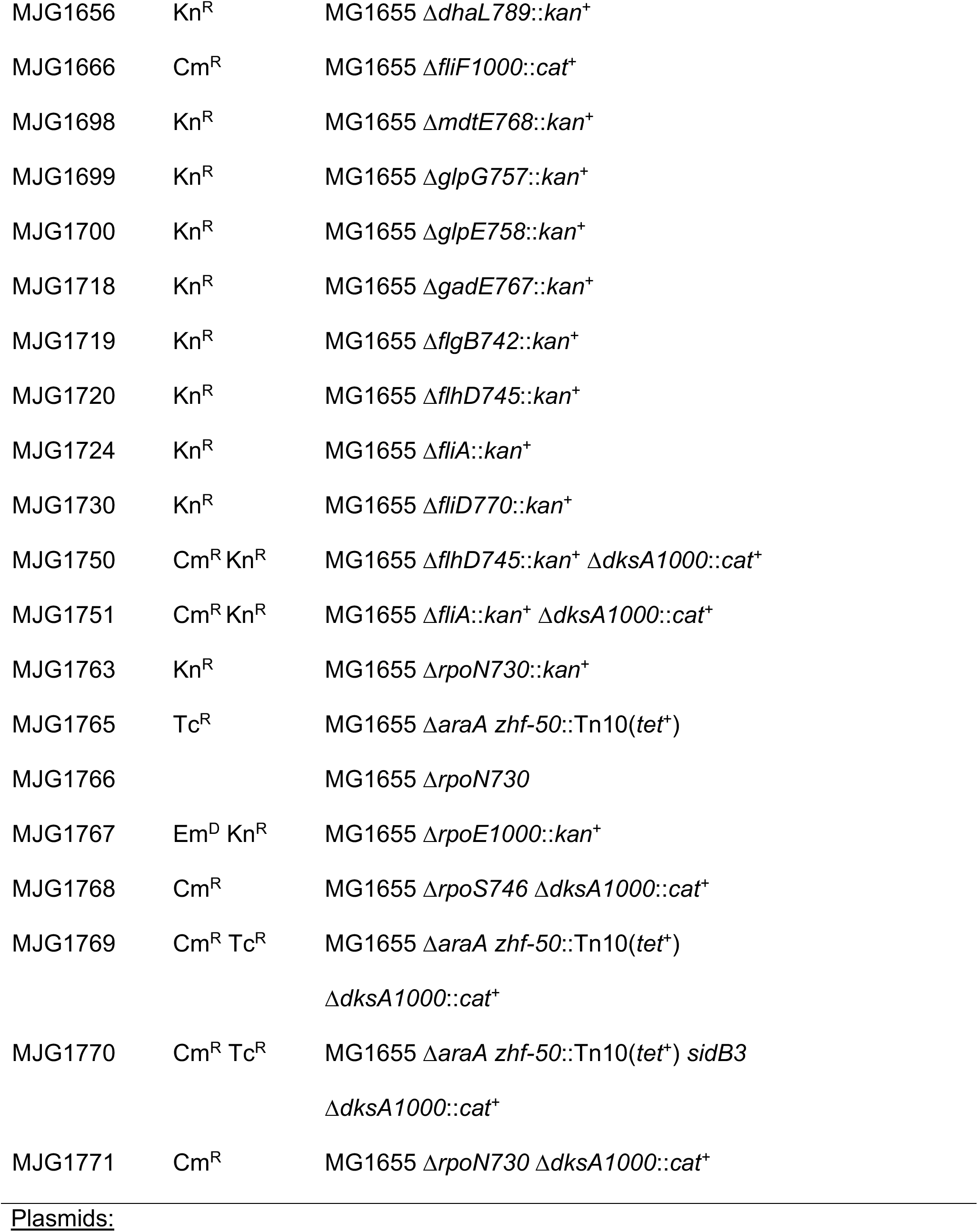
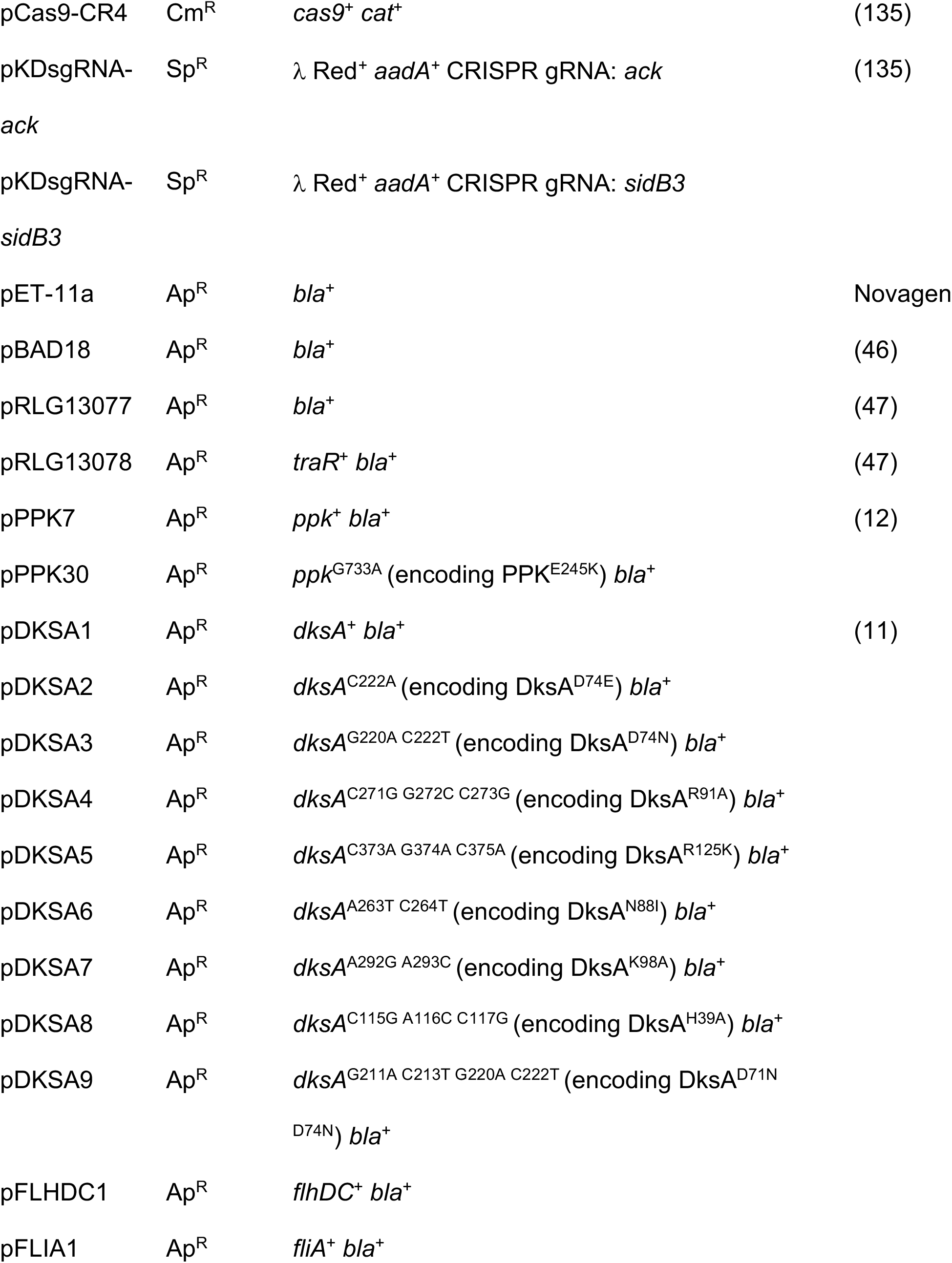
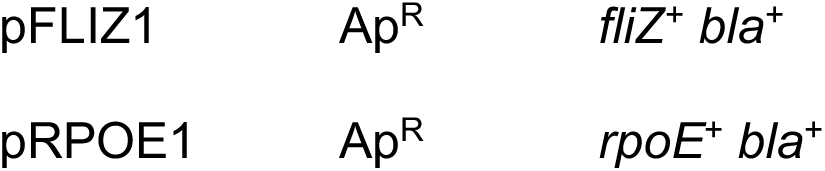
Strains and plasmids used in this study. Unless otherwise indicated, all strains and plasmids were generated in the course of this work. Abbreviations: Ap^R^, ampicillin resistance; Cm^R^, chloramphenicol resistance; Em^D^, erythromycin dependence; Kn^R^, kanamycin resistance; Rif^R^, rifampicin resistance; Sm^R^, streptomycin resistance; Sp^R^, spectinomycin resistance; Tc^R^, tetracycline resistance.

### Databases and primer design

I obtained gene and protein sequences and metabolic pathway information from the Integrated Microbial Genomes database (125) and from EcoCyc (72), and obtained information about *E. coli* regulatory networks from RegulonDB (126). I designed PCR and sequencing primers with Web Primer (www.candidagenome.org/cgi-bin/compute/web-primer), and mutagenic primers with PrimerX (www.bioinformatics.org/primerx/index.htm). I designed all primers used for qPCR with Primer Quest (www.idtdna.com; parameter set “qPCR 2 primers intercalating dyes” for qRT-PCR primer design), and tested each primer pair to confirm specificity and amplification efficiencies of close to 1. These primers are listed in Table 2.

**TABLE 2.**
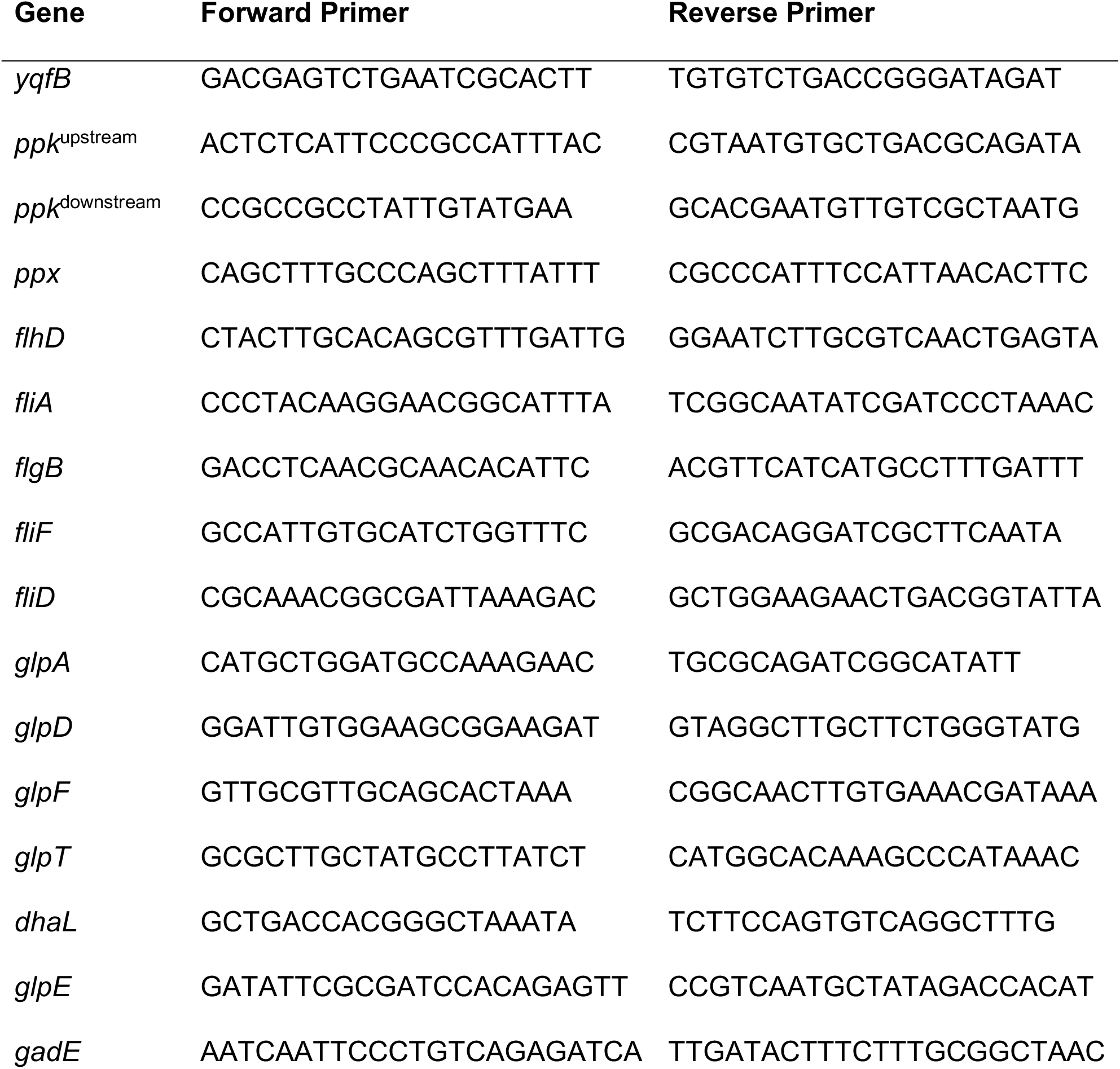
Primers used for quantitative RT-PCR

### Strain construction

Unless otherwise indicated, all *E. coli* strains were derivatives of wild-type strain MG1655 (F^-^, *rph-1 ilvG^-^ rfb-50*) (127). I confirmed all chromosomal mutations by PCR. Strain MC4100 Δ*dnaK52* (F^-^, λ^-^, e14^-^, [*araD139*]B/r Δ(*argF-lac*)*169 flhD5301* Δ(*fruK-yeiR*)*725*(*fruA25*) *relA1 rpsL150*(Sm^R^) *rbsR22* Δ(*fimB-fimE*)*632*(::IS1) *deoC1* Δ*dnaK52*::*cat*^+^)(60, 128) was a gift from Ursula Jakob (University of Michigan), and strain CAG18450 (*zhf-50*::Tn10(*tet*^+^))(129, 130) was a gift from Chuck Turnbough (University of Alabama at Birmingham).

I used P1*vir* transduction (11, 131) to move the Δ*greA788*::*kan*^+^ allele from the Keio collection (45) into MG1655 and MJG1419 (Δ*dksA1000*::*cat*^+^)(11), generating strains MJG1559 (Δ*greA788*::*kan*^+^) and MJG1561 (Δ*dksA1000*::*cat*^+ Δ^*greA788*::*kan*^+^).

I generated strain MJG0639 (Δ*araA*::*cat*^+^ Δ*ppk-749*) by P1*vir* transduction of the Δ*araA*::*cat^+^* allele from strain MT555 (Δ*malE ΔhsdR ΔaraA*::*cat^+^*) (a gift from James Bardwell, University of Michigan) into MJG0224 (Δ*ppk-749*)(12). I then resolved the chloramphenicol resistance cassette in this strain (109), yielding strain MJG0640 (Δ*araA* Δ*ppk-749*). I used P1*vir* transduction to move the *ΔdnaK52*::*cat*^+^ allele from strain MC4100 *ΔdnaK52*::*cat*^+^ into strains MJG0621 (Δ*araA*)(34) and MJG0640 (Δ*araA* Δ*ppk-749*), yielding strains MJG1449 (Δ*araA ΔdnaK52*::*cat*^+^) and MJG1461 (Δ*araA* Δ*ppk-749 ΔdnaK52*::*cat*^+^). Unless otherwise indicated, I grew all Δ*dnaK52*::*cat*^+^ strains at 30°C.

I moved the Δ*glpA721*::*kan*^+^, Δ*glpD759*::*kan*^+^, Δ*glpF786*::*kan*^+^, Δ*glpT720*::*kan*^+^, Δ*mdtE768*::*kan*^+^, Δ*glpG757*::*kan*^+^, Δ*glpE758*::*kan*^+^, Δ*gadE767*::*kan*^+^, Δ*flgB742*::*kan*^+^, Δ*flhD745*::*kan*^+^, Δ*fliA*::*kan*^+^ (no allele number provided in the Coli Genetic Stock Center database; cgsc.biology.yale.edu), and Δ*fliD770*::*kan*^+^ alleles from the Keio collection by P1*vir* transduction into MG1655, generating strains MJG1651 (Δ*glpA721*::*kan*^+^), MJG1652 (Δ*glpD759*::*kan*^+^), MJG1653 (Δ*glpF786*::*kan*^+^), MJG1654 (Δ*glpT720*::*kan*^+^), MJG1698 (Δ*mdtE768*::*kan*^+^), MJG1699 (Δ*glpG757*::*kan*^+^), MJG1700 (Δ*glpE758*::*kan*^+^), MJG1718 (Δ*gadE767*::*kan*^+^), MJG1719 (Δ*flgB742*::*kan*^+^), MJG1720 (Δ*flhD745*::*kan*^+^), MJG1724 (Δ*fliA*::*kan*^+^), and MJG1730 (Δ*fliD770*::*kan*^+^). I used P1*vir* transduction to move the Δ*dksA1000*::*cat*^+^ allele from MJG1419 into MJG1720 (Δ*flhD745*::*kan*^+^) and MJG1724 (Δ*fliA*::*kan*^+^), yielding strains MJG1750 (Δ*flhD745*::*kan*^+^ Δ*dksA1000*::*cat*^+^) and MJG1751 (Δ*fliA*::*kan*^+^ Δ*dksA1000*::*cat*^+^).

I replaced the *fliF* gene of strain MG1655 with a pKD3-derived chloramphenicol resistance cassette by recombineering (109), using primers 5’ GTT CCA CTT TGC CAA TAA CGC CGT CCA TAA TCA GCC ACG AGG TGC GCG ATG GTG TAG GCT GGA GCT GCT TC 3’ and 5’ TCG CCA ATG GTC ATC AGC AGG ATG ACG CTT TTA TCG GTG CCT GTC AGG TTA CAT ATG AAT ATC CTC CTT AG 3’ and yielding strain MJG1666 (Δ*fliF1000*::*cat*^+^). I amplified the Δ*dhaL789*::*kan*^+^ allele from the Keio collection with primers 5’ TTC TCA ATC ACC TTA CTG 3’ and 5’ CAA CTA TCA CTC ATT AAC 3’ and used the resulting product as a recombineering substrate in MG1655, yielding strain MJG1656 (Δ*dhaL789*::*kan*^+^).

I used oligo-directed recombineering (132) to construct a chromosomal *rpoB3443* mutation (83-85, 133, 134) using the mutagenic primer 5’ AAG CCT GCA CGT TCA CGG GTC AGA CCG CCG GGA CCG GGG GCA GAG ATA CGA CGT TTG TGC GTA ATC TCA GAC A 3’, which contained four 5’ phosphorothiorate linkages. This primer generates an *rpoB*^C1593T, A1596C, T1598C, C1602T, A1605C^ allele, encoding RpoB^L533P^, with silent mutations in four codons adjacent to codon 533 to avoid mismatch repair. I transformed MG1655 with pKD46 (109), induced expression of *λ* Red recombinase, electroporated with 250 pmol of mutagenic primer, and selected recombinant colonies at 37°C on LB plates containing rifampicin. I confirmed the sequence of *rpoB* alleles by PCR amplification and Sanger sequencing of a fragment of *rpoB* with primers 5’ GAT GTT ATG AAA AAG CTC 3’ and 5’ CTG GGT GGA TAC GTC CAT 3’. After curing pKD46 by growth at 37°C, this yielded strain MJG1377 (*rpoB3443*).

I used P1*vir* transduction to move the Δ*rsd-766*::*kan*^+^ and Δ*rpoN730*::*kan*^+^ alleles from the Keio collection into MG1655, generating strains MJG1376 (Δ*rsd-766*::*kan*^+^) and MJG1763 (Δ*rpoN730*::*kan*^+^), then resolved the kanamycin resistance cassette in MJG1763 (109), yielding strain MJG1766 (Δ*rpoN730*). I used P1*vir* transduction to move the Δ*dksA1000*::*cat*^+^ allele from MJG1419 into MJG0344 (Δ*rpoS746*)(11) and MJG1766 (Δ*rpoN730*), yielding strains MJG1768 (Δ*rpoS746* Δ*dksA1000*::*cat*^+^) and MJG1771 (Δ*rpoN730* Δ*dksA1000*::*cat*^+^).

I used Cas9-assisted recombineering (135) to construct a chromosomal *sidB3* (60) mutation of strain CAG18450 (*zhf-50*::Tn10(*tet*^+^))(129, 130) using the mutagenic primer 5’ ACT CTC ATC CAG GGT TCT CTG CTT AAT AGC GGG AAT TCG GCC TCG ATG GCA GCA CGC AAT TTT TTC ATC GCG T 3’, which contained four 5’ phosphorothiorate linkages. This primer mutates the TAA stop codon of *rpoH* to a GAA glutamate codon (*rpoH*^T853G^), and makes silent mutations in 4 flanking codons to avoid mismatch repair. I confirmed the sequence of *rpoH* alleles by PCR amplification and Sanger sequencing of a fragment of *rpoH* and its 3’ downstream sequence with primers 5’ TGT CTT CCG ACG ACG ATT 3’ and 5’ GTA ACG CTT TAC CCT TTA 3’. Due to the temperature sensitivity of a *sidB3* mutant (60), it was not possible to cure the pKDsgRNA-*sidB3* protospacer targeting plasmid (see below) by growth at 42°C (135), so I used P1*vir* transduction to move *sidB3* and the 60% linked *zhf-50* tetracycline resistance marker (transposon Tn10 in the intergenic region between *ugpB* and *ilvF*)(129, 130) from the resulting strain into MJG0621 (Δ*araA*) to generate strain MJG1448 (Δ*araA zhf-50*::Tn10(*tet*^+^) *sidB3*). I used P1*vir* transduction to move *zhf-50* from CAG18450 to MJG0621 (Δ*araA*) to generate strain MJG1765 (Δ*araA zhf-50*::Tn10(*tet*^+^)). I also used P1*vir* transduction to move the *dksA1000*::*cat*^+^ allele from MJG1419 into MJG1765 (Δ*araA zhf-50*::Tn10(*tet*^+^)) and MJG1448 (Δ*araA zhf-50*::Tn10(*tet*^+^) *sidB3*), yielding strains MJG1769 (Δ*araA zhf-50*::Tn10(*tet*^+^) Δ*dksA1000*::*cat*^+^) and MJG1770 (Δ*araA zhf-50*::Tn10(*tet*^+^) *sidB3* Δ*dksA1000*::*cat*^+^). I maintained all *sidB3* strains at 30°C.

I replaced the *rpoE* gene of strain MG1655 with a pKD4-derived kanamycin resistance cassette by recombineering (109), using primers 5’ GCG TTT CGA TAG CGC GTG GAA ATT TGG TTT GGG GAG ACT TTA CCT CGG ATG GTG TAG GCT GGA GCT GCT TC 3’ and 5’ GTT GTT CTT TCT GCA TGC CTA ATA CCC TTA TCC AGT ATC CCG CTA TCG TCA CAT ATG AAT ATC CTC CTT AG 3’. This resulted in strain MJG1767 (Δ*rpoE1000*::*kan*^+^). I made multiple attempts to use P1*vir* transduction to move the Δ*dksA1000*::*cat*^+^ allele from MJG1419 into MJG1767 (Δ*rpoE1000*::*kan*^+^), without success. I constructed, maintained, and tested all *rpoE* mutant strains in media containing 10 µg ml^-1^ erythromycin (104).

### Plasmid construction

Plasmids pRLG13077 and pRLG13078 (47) were a gift from Richard Gourse and Saumya Gopalkrishnan (University of Wisconsin-Madison). pCas9-CR4 and pKDsgRNA-*ack* were gifts from Kristala Prather (Addgene plasmid # 62655; http://n2t.net/addgene:62655; RRID:Addgene_62655 and Addgene plasmid # 62654; http://n2t.net/addgene:62654; RRID:Addgene_62654)(135).

I replaced the sgRNA targeting sequence of plasmid pKDsgRNA-*ack* by round-the-horn cloning (135) with primer 5’ TAG CGG AAA TTA CGC TTC AAG TTT TAG AGC TAG AAA TAG CAA G 3’, yielding plasmid pKDsgRNA-*sidB3*.

I used the QuikChange^™^ site-directed mutagenesis method (Agilent Technologies), modified to use a single primer and 35 cycles of amplification, to mutate the pBAD18-derived (46) *ppk*^+^ plasmid pPPK7 (136) with primer 5’ GAA GCC AGC CTG ATG AAG TTG ATG TCT TCC 3’ to generate plasmid pPPK30, containing a *ppk*^G733A^ allele encoding PPK^E245K^. I used the same procedure to mutate the pBAD18-derived *dksA*^+^ plasmid pDKSA1 (11) as follows: I used primer 5’ CAA CTT CCC GGA CCC GGT AGA ACG TGC AGC CCA GGA AGA AG 3’ to generate plasmid pDKSA2 (*dksA*^C222A^, encoding DksA^D74E^); I used primer 5’ CAA CTT CCC GGA CCC GGT AAA TCG TGC AGC CCA GGA AGA AG 3’ to generate plasmid pDKSA3 (*dksA*^G220A C222T^, encoding DksA^D74N^); I used primer 5’ CTC GAA CTG CGT AAC CGC GAT GCG GAG CGT AAG CTG ATC AAA AAG 3’ to generate plasmid pDKSA4 (*dksA*^C271G G272C C273G^, encoding DksA^R91A^); I used primer 5’ GTT GAA ATT GGT ATT CGC AAA CTG GAA GCG CGC CCG AC 3’ to generate plasmid pDKSA5 (*dksA*^C373A G374A C375A^, encoding DksA^R125K^), I used primer 5’ CAG CCT CGA ACT GCG TAT TCG CGA TCG CGA GCG TAA G 3’ to generate plasmid pDKSA6 (*dksA*^A263T C264T^, encoding DksA^N88I^), I used primer 5’ CGT AAG CTG ATC AAA GCG ATC GAG AAG ACG C 3’ to generate plasmid pDKSA7 (*dksA*^A292G A293C^, encoding DksA^K98A^), and I used primer 5’ GAA TGA AGC CCA GCT GGC GGC GTT CCG TCG TAT TCT GGA AG 3’ to generate plasmid pDKSA8 (*dksA*^C115G A116C C117G^, encoding DksA^H39A^). Finally, I mutated plasmid pDKSA3 with primer 5’ GCA GCC AAC TTC CCG AAT CCG GTA AAT CGT GCA G 3’ to generate plasmid pDKSA9 (*dksA*^G211A C213T G220A C222T^, encoding DksA^D71N D74N^).

I amplified the *flhDC* coding sequence (932 bp) from *E. coli* MG1655 genomic DNA with primers 5’ TTC GAA TTC AAG GAG ATA TAC ATA TGC ATA CCT CCG AGT TGC TGA AA 3’ and 5’ CTT AAG CTT TTA AAC AGC CTG TAC TCT CTG TTC A 3’, incorporating an AAG GAG ATA TAC AT ribosome binding site (RBS) sequence derived from pET-11a (Novagen), and cloned it into the *Eco*RI and *Hin*dIII sites of plasmid pBAD18 (46) to generate plasmid pFLHDC1. Similarly, I amplified the *fliA* and *fliZ* coding sequences (720 and 552 bp, respectively) with primers 5’ ACC GGT ACC AAG GAG ATA TAC ATA TGA ATT CAC TCT ATA CCG CTG AAG G 3’ and 5’ CTT AAG CTT TTA TAA CTT ACC CAG TTT AGT GCG T 3’ or 5’ TTC GAA TTC AAG GAG ATA TAC ATA TGA TGG TGC AGC ACC TGA AAA 3’ and 5’ CTT AAG CTT TTA ATA TAT ATC AGA AGA AGG CAG GCT GG 3’, generating products with the above RBS and, in the case of *fliA*, changing the native GTG start codon to an ATG. I cloned these products into the *Kpn*I and *Hin*dIII or *Eco*RI and *Hin*dIII sites of pBAD18 to generate plasmids pFLIA1 and pFLIZ1. I amplified *rpoE* (576 bp) from *E. coli* MG1655 genomic DNA with primer pair 5’ TTT GAA TTC AAG GAG ATA TAC ATA TGA GCG AGC AGT TAA CGG A 3’ and 5’ AGA TCT AGA TCA ACG CCT GAT AAG CGG TT 3’, incorporating the pET-11a RBS, and cloned the resulting product into the *Eco*RI and *Xba*I sites of plasmid pBAD18 (46) to generate plasmid pRPOE1.

### *In vivo* polyphosphate assay

To induce polyP synthesis by nutrient limitation (10, 11, 34), I grew *E. coli* strains in 10 ml LB at 30°C or 37°C with shaking (200 rpm) to A_600_=0.2-0.4. I harvested samples (1 ml; 50-100 µg of total protein) for polyP measurements as described below, and then harvested 5 ml of each culture by centrifugation (5 min at 4,696 x g at room temperature), resuspended them in 5 ml phosphate-buffered saline (PBS) to rinse, then re-centrifuged and resuspended them in 5 ml MOPS minimal medium (Teknova)(137) containing 4 g l^-1^ glucose, 0.1 mM K_2_HPO_4_, and 0.1 mM uracil. I incubated these cultures for 2 hours at 30°C or 37°C with shaking, then collected additional samples for polyP measurements. Where indicated, I added arabinose (0.125 or 2 g l^-1^), IPTG (1 mM), or erythromycin (10 µg ml^-1^) to both the LB and the MOPS medium.

I measured intracellular polyP levels as previously described (138). Briefly, I harvested samples of bacterial cultures by centrifugation, resuspended them in 250 µl of 4 M guanidine isothiocyanate, 50 mM Tris-HCl (pH 7), lysed them by incubation for 10 min at 95°C, and immediately froze them at -80°C. After thawing at room temperature, I determined protein concentrations by Bradford assay (Bio-Rad) of 5 µl aliquots of each sample. I added 250 µl of 95% ethanol to the remaining sample and applied the resulting mixture to an EconoSpin silica spin column (Epoch Life Science), rinsed with 750 µl 5 mM Tris-HCl, pH 7.5, 50 mM NaCl, 5 mM EDTA, 50% ethanol, and eluted with 150 µl 50 mM Tris-HCl, pH 8. I brought the eluate to 20 mM Tris-HCl, pH 7.5, 5 mM MgCl_2_, 50 mM ammonium acetate with 1 µg of PPX1 exopolyphosphatase from *Saccharomyces cereviseae* (139) in a final volume of 200 µl and incubated for 15 min at 37°C, then measured the resulting polyP-derived orthophosphate using a colorimetric assay (140) and normalized to total protein content. For all figures, I reported polyP concentrations in terms of individual phosphate monomers.

### Quantitative RT-PCR

I stressed *E. coli* strains by nutrient limitation as described above. At the indicated time points, I harvested 1 ml of cells by centrifugation and resuspended in RNA*later* (ThermoFisher) for storage at -20°C. I extracted RNA using the RiboPure™ RNA Purification Kit for bacteria (Ambion) following the manufacturer’s instructions, including the optional DNAse treatment to remove contaminating genomic DNA. I used the SuperScript^TM^ IV VILO^TM^ kit (ThermoFisher) to reverse transcribe cDNA from mRNA, following the manufacturer’s instructions, including a no-RT control for each reaction. I calculated changes in gene expression using the 2^-ΔΔCt^ method (141), with *yqfB* as an internal expression control. (Since both nutrient limitation and mutation of *dksA* were expected to change the expression of ribosomal RNA genes (5), which are common housekeeping genes used to normalize qRT-PCR data in bacteria (142), I used *yqfB* instead, the expression of which did not change under the conditions tested here (Supplemental Dataset S1).)

### RNA sequencing

I grew the indicated *E. coli* strains as described above for induction of polyP synthesis by nutrient limitation, and purified RNA from 2 ml of cells harvested immediately before and 5 min after stress using the RiboPure™-Bacteria Kit (Ambion), according to the manufacturer’s instructions. mRNA-sequencing was performed by the UAB Heflin Center for Genomic Sciences, using an Illumina NextSeq 500 as described by the manufacturer (Illumina, Inc). Briefly, the quality of the total RNA was assessed using a Agilent 2100 Bioanalyzer and RNA with a RNA Integrity Number (RIN) of 7.0 or above was used for sequencing library preparation. The Agilent SureSelect Strand Specific mRNA library kit was used according to the manufacturer’s instructions (Agilent). Library construction began with ribosome reduction with the RiboMinus protocol for Gram-negative and Gram-positive bacteria as described by the manufacturer (Life Technologies). The resulting mRNA was randomly fragmented with cations and heat, followed by first strand synthesis using random primers with inclusion of Actinomycin D (2.4 ng µl^-1^ final concentration). Second strand cDNA production was done with standard techniques, the ends of the resulting cDNA were made blunt, A-tailed, and adaptors were ligated for amplification and indexing to allow for multiplexing during sequencing. The cDNA libraries were quantitated using qPCR in a Roche LightCycler 480 with the Kapa Biosystems kit for Illumina library quantitation (Kapa Biosystems) prior to cluster generation. Cluster generation was performed according to the manufacturers recommendations for onboard clustering (Illumina). Approximately 20 million single end 75 bp reads were generated per sample, and I analyzed the resulting data using the Rockhopper 2 software package (143) and Morpheus (https://software.broadinstitute.org/morpheus).

### Statistical analyses

I used GraphPad Prism version 8.3.1 for Macintosh (GraphPad Software) to perform statistical analyses, including two-way repeated measures ANOVA with Holm-Sidak’s multiple comparison tests. Repeated measures ANOVA cannot handle missing values, so data sets with samples having different *n* numbers (*e.g.* Fig. 2) were analyzed with a mixed model which uses a compound symmetry covariance matrix and is fit using Restricted Maximum Likelihood (REML) (without Geisser-Greenhouse correction).

### Data availability

All strains and plasmids generated in the course of this work are available from the author upon request. RNA sequencing data have been deposited in NCBI’s Gene Expression Omnibus (144) and are accessible through GEO Series accession number GSE144816.

## ACKNOWLEDGEMENTS

This project was supported by University of Alabama at Birmingham Department of Microbiology startup funds and NIH grant R35GM124590. The author has no conflicts of interest to declare, and would like to thank Dr. Avishek Mitra (UAB Department of Microbiology) and Drs. Saumya Gopalkrishnan, Rick Gourse, and Wilma Ross (University of Wisconsin – Madison) for helpful discussions of the data presented here.

